# Structural elements required for the efficient loading and activation of HELB on RPA-coated single-stranded DNA

**DOI:** 10.64898/2026.07.16.738708

**Authors:** Oliver J. Wilkinson, Silvia Hormeño, Clara Aicart-Ramos, Ayush Mistry, Edwin Antony, Fernando Moreno-Herrero, Mark S. Dillingham

**Author notes:** Corresponding authors: Mark S. Dillingham and Fernando Moreno-Herrero. These authors contributed equally (Oliver J. Wilkinson, Silvia Hormeño). Classification: Biological Sciences - Biochemistry.

## Abstract

HELB is a human helicase involved in DNA repair and replication that interacts physically with the single-stranded DNA binding protein RPA. ATP-dependent translocation of HELB along ssDNA results in the active displacement of RPA molecules and the formation of ssDNA loops, suggesting that HELB contains at least two DNA binding sites. In this work, we investigated the role of HELB-specific structural elements in facilitating interactions between HELB and RPA-coated DNA. We show that a predicted OB-fold in the N-terminal region of the protein is important both for loop extrusion and RPA displacement. We confirm that a HELB-specific-motif within the RecA-like helicase/translocase domains is critical for binding RPA in solution but that, once HELB is bound to ssDNA, is dispensable for RPA displacement. We propose a model for RPA displacement in which both structural elements play important roles in the recruitment and activation of HELB at RPA-ssDNA filaments.

**SIGNIFICANCE STATEMENT:** Single-stranded DNA generated during replication and repair is rapidly coated by replication protein A (RPA), creating a protected filament that must nevertheless remain accessible to DNA-processing enzymes. We show that the human DNA helicase HELB uses two specialised structural elements to overcome this problem. A HELB-specific RPA-binding motif promotes recruitment, whereas a predicted OB domain enables DNA looping and efficient RPA displacement. Removing these elements modestly enhances activity on naked ssDNA while impairing loading and activation on RPA-coated DNA. This reveals how helicase accessory domains restrain inappropriate motor activity while targeting the intended nucleoprotein substrate. Human HELB variants linked to reproductive ageing map to these regulatory regions, suggesting a connection between impaired RPA dynamics and reproductive health.

## INTRODUCTION

Human HELB protein is a Superfamily 1B (SF1B) helicase with reported roles in multiple DNA replication and repair pathways [1, 2]. These include both the inhibition [3] and stimulation [4] of homology-dependent DNA repair, the initiation of chromosomal DNA replication [5], recovery from replicative stress [6], and the resolution of secondary structures formed by CGG nucleotide repeats [7]. Single nucleotide polymorphisms in the HELB gene are associated with fertility traits in mammals [8–11]. Given the established links between oocyte health and DNA repair pathways this might reflect an important role for HELB in meiotic recombination and/or recovery from DNA damage in oocytes [12].

In previous work, we showed that HELB efficiently and specifically displaces human RPA from ssDNA. RPA is the primary single-stranded DNA-binding protein in eukaryotes and plays a crucial role in DNA replication, recombination and repair [13]. It binds ssDNA rapidly and with very high affinity, effectively coating and protecting exposed ssDNA from degradation. Beyond this protective function, RPA also mediates a variety of downstream processes involving ssDNA and, in this context, RPA interacts with numerous proteins implicated in DNA metabolism including many helicases [14–16]. RPA is a dynamic heterotrimer composed of RPA70, RPA32 and RPA14 subunits and including six oligonucleotide binding (OB) folds forming DNA-binding domains (DBDs) linked by flexible regions [17–20]. Protein interactions with RPA are commonly mediated through the N-terminal domain of RPA70 (RPA70N) and/or the C-terminal winged-helix of RPA32. Because RPA-ssDNA filaments are a crucial intermediate in all pathways in which HELB is reported to operate, we argued that the RPA-remodelling activity might underpin all cellular functions of HELB [21]. The primary aim of the present study is to investigate the mechanistic basis for the specific targeting of RPA-ssDNA filaments by HELB.

HELB is composed of three functional domains: an N-terminal domain (NTD) of undefined structure, a helicase domain containing the seven conserved SF1B motifs which form the primary ssDNA binding/translocation site [2, 22], and a C-terminal domain (CTD) that contains cyclin-dependent kinase (CDK) phosphorylation sites and a subcellular localization domain. The extended N-terminal region of HELB is not conserved in otherwise related proteins such as RecD2 [23], and its role is not yet fully understood. One study shows that it is important for interaction with Cdc45, a component of the replicative helicase CMG [24]. Moreover, in our previous paper we suggested that the NTD might contain a secondary single-stranded DNA binding site, in part because HELB is a monomeric protein that is able to loop ssDNA during translocation [21]. Modeling of HELB predicts the presence of an oligonucleotide/oligosaccharide binding (OB) fold in the first 115 amino acids of the NTD (**Figure 1A**), which is very close to the site of a human polymorphism (D117Y) associated with premature ovarian insufficiency and early age of natural menopause [11]. In the work presented below, we shall refer to this region of the protein as the OB domain, but it should be noted that this is a prediction; there is no experimentally validated structural data available for HELB except for a peptide from the HSM domain [25].

**Figure 1:**
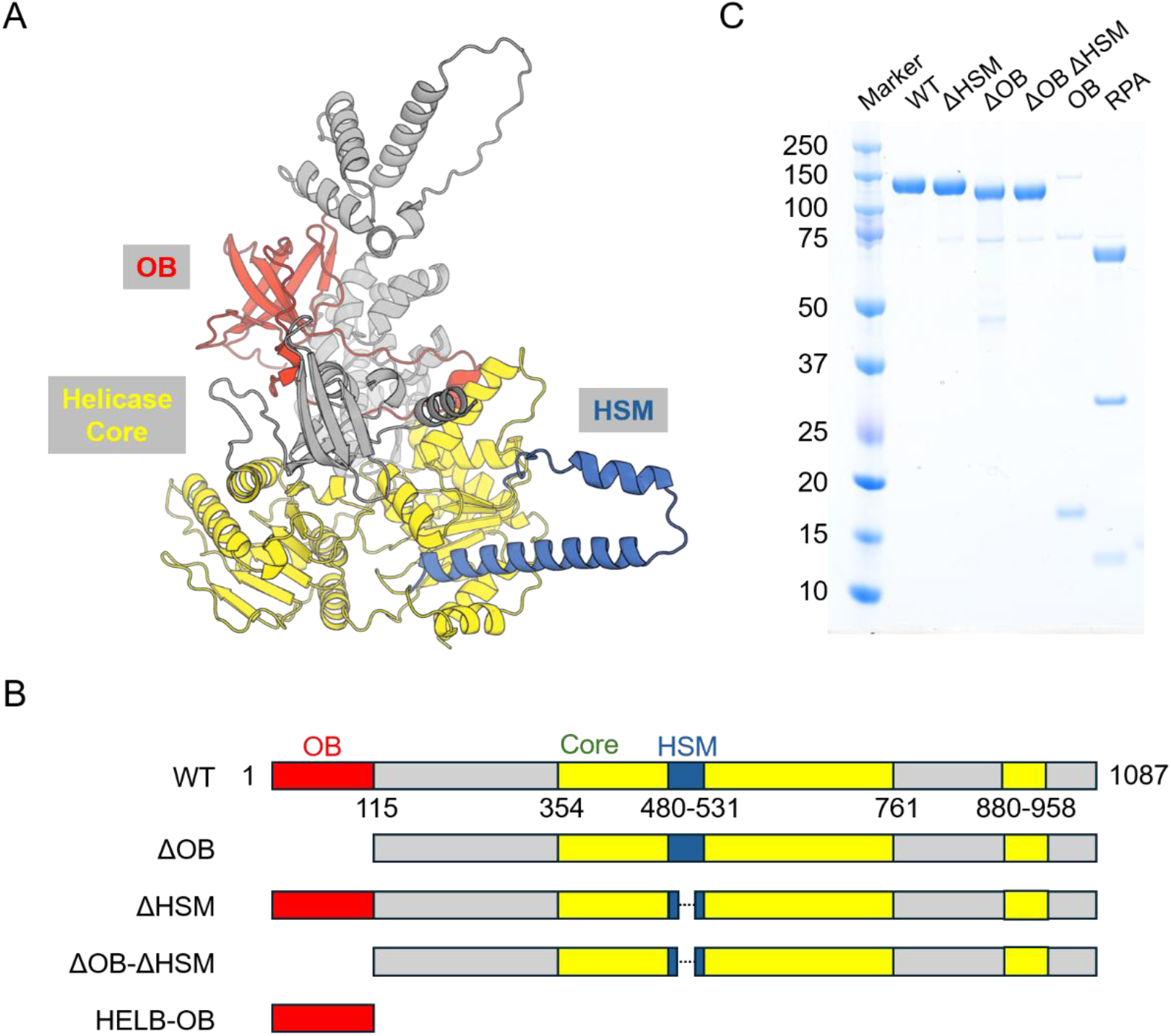
Domain organisation of HELB. (A) The structure shown is an AlphaFold3 model of human HELB colour-coded according to the primary structure diagram below [27]. Model metrics were pTM, 0.75; predicted disordered fraction, 0.25; atom-level pLDDT mean, 71; atom-level pLDDT median 83, indicating a confidently modelled core with lower confidence in flexible/disordered regions. The figure highlights an OB-domain (red), the helicase core domains (yellow) and the HELB Specific Motif (HSM) (blue). Disordered loops with low confidence scores have been removed from the model for clarity. (B) The primary structure diagram highlights the domain organisation of wild type HELB and the design of the variant proteins used in this study. (C) Purified proteins used in this study analysed by SDS-PAGE.

In addition to providing the motor activity that drives ssDNA translocation and RPA displacement, the central helicase domain of HELB has been reported to interact with the RPA70N domain through an acidic motif located between helicase motif I and motif Ia [6]. This ‘HELB-specific motif’ (HSM) comprises residues 493-522 and is unique to HELB family members [6, 23] implying that its interaction with RPA may relate to the specific function of this helicase. A recently solved structure of the HELB-HSM peptide (residues 496-519 of HELB) bound to RPA70N showed that the motif consists of a 4- turn α-helix followed by a β-turn and a 3_10_ helix, which fits into the shallow basic groove of RPA70N with HELB residues E496, E499 and D506 playing a central role in stabilizing the interaction [25]. Importantly, some genetic variants of HELB implicated in reproductive ageing traits (D506G and E522D) map to this interface within RPA, suggesting that failure of HELB to interact with RPA may impact human health [1, 26].

In a previous study, we found that HELB and RPA form a complex on ssDNA in ensemble assays and that human RPA specifically stimulates the ATPase activity of HELB [21]. We also demonstrated that HELB functions as an efficient ssDNA translocase with the ability to extrude ssDNA loops but limited unwinding capacity, highlighting that RPA-ssDNA is its primary substrate. Furthermore, we observed that HELB’s processive translocation is accompanied by the displacement of RPA from ssDNA. These findings provided new mechanistic insight into HELB’s function but also presented a paradox: how can RPA (an exceptionally tight ssDNA binding protein) recruit and activate HELB to act on the same underlying substrate. Moreover, the mechanistic basis for the observed ssDNA looping activity was not resolved. In this work, we investigated the structural basis for the recruitment and activation of HELB on RPA-ssDNA filaments. By comparing the activity of wild-type HELB with mutants lacking the OB-fold and/or the HELB specific motif we show that both structural elements are dispensable for HELB activity on naked ssDNA but play crucial roles in the recruitment and activation of HELB on RPA-ssDNA filaments.

## RESULTS

### Design and purification of HELB variant proteins

To guide the design of HELB variant proteins we used Alphafold3 to model the structure (**Figure 1A**) [27]. The model metrics (pTM, 0.75; predicted disordered fraction, 0.25; atom-level pLDDT mean, 71; atom-level pLDDT median 83) indicate a confidently modelled core with lower certainty in flexible/disordered regions. To better understand how HELB clears RPA from ssDNA during translocation, and whether this process is mediated by specific interactions between HELB and RPA, we engineered HELB variants targeting two regions of the structure that might play a key role in the process. One variant, which we call ΔOB, contains a deletion of the first 117 amino acids which removes a predicted OB-fold (mean pLDDT = 71; **Figure 1B, red domain**). A second variant protein, ΔHSM, was engineered which replaces a region of the helicase core spanning residues 496-531 with a short linker sequence. This removes critical residues from a region of the protein called the HELB- specific motif (HSM) which has been implicated in RPA-interaction in multiple studies (**Figure 1B, blue domain**) [6, 25, 26]. We also produced a ΔOB-ΔHSM variant containing both deletions, as well as a construct named HELB-OB which only contains the predicted N-terminal OB fold (residues 1-115). Literature reviews and primary structure analyses identified potential RPA interaction motifs in the OB and HSM domains that might bind the N-terminal region of RPA70 (**SFigure 1**). Therefore, our deletion constructs also have the effect of removing either (or both) of these putative RPA70N binding motifs from the HELB protein. All constructs were expressed in insect cells and purified to homogeneity as described in the Methods section (**Figure 1C**).

### HELB variants retaining the core domains display wild-type or better than wild-type ssDNA binding activity

We first tested properties of HELB that were independent of its interaction with RPA by using DNA binding and ATP hydrolysis assays to study the interaction between the HELB variants, naked ssDNA and ATP. Wild type HELB shifted a 25mer ssDNA oligonucleotide to form two distinct complexes in a native gel, likely representing complexes with one or two bound HELB molecules (**Figure 2A**). The three variant proteins (ΔHSM, ΔOB and ΔOB-ΔHSM) behaved similarly to wild type in this assay both quantitatively and qualitatively, and in a reproducible manner (**Figure 2A**; additional example in **SFigure 2A**). We also reproducibly detected very weak (micromolar range) binding to DNA using preparations of purified HELB-OB domain in this assay (**Figure 2B** and **SFigure 2B**) supporting the idea that this region of the protein does indeed form a *bona fide* OB domain that has been shown to interact with DNA in many other proteins (**Figure 1A**).

**Figure 2:**
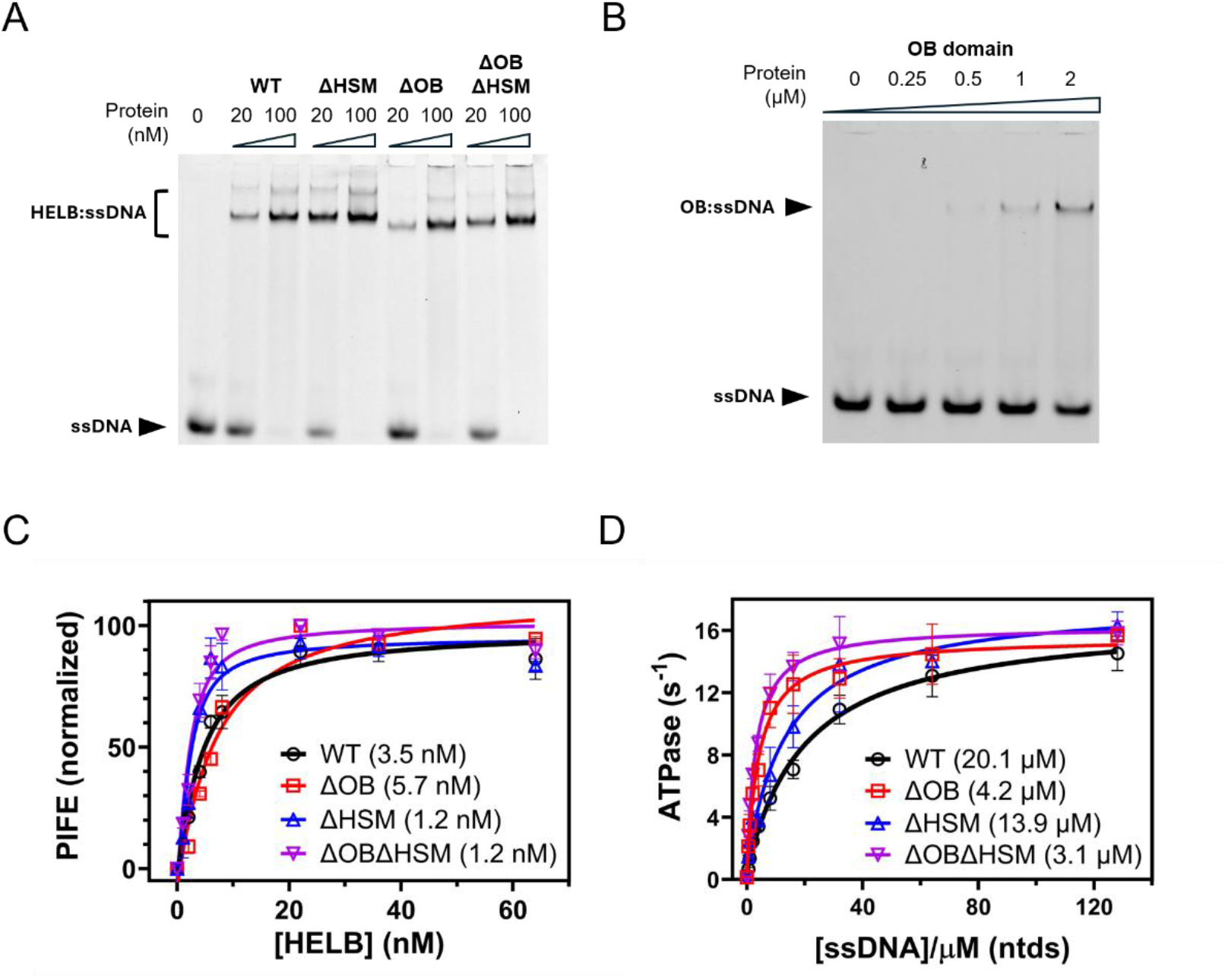
HELB variants retaining the core domains display wild-type or better than wild-type ssDNA binding activity. (A) EMSA assays showing binding of a 25mer ssDNA oligonucleotide (5 nM) by wild-type and variant HELB proteins at the concentrations indicated. A further example of this assay is shown in Figure S2A to demonstrate reproducibility. (B) EMSA assays showing binding of a 25mer ssDNA oligonucleotide (5 nM) by HELB-OB domain at the concentrations indicated. Note that the apparent binding is very weak. A further example of this assay is shown in Figure S2B to demonstrate reproducibility. (C) Protein-induced fluorescence enhancement assay showing binding of a 3′-Cy3-labelled 40mer ssDNA oligonucleotide by wild type and variant HELB proteins. The K_d_ values obtained from the fits are shown. Note that the binding of variant proteins to ssDNA is comparable to, or slightly better, than wild type. (D) ssDNA-dependent ATPase activity of wild type and variant HELB proteins. The K_DNA_ values obtained from the fits are shown. Note that the apparent binding of the variant proteins to ssDNA is better than wild type.

We next used protein-induced fluorescence enhancement (PIFE) assays to generate binding isotherms for the wild type and variant proteins with ssDNA. These were fit to a one-site binding equation to yield Kd (see Methods for details). Wild type HELB interacted with a Kd value of 3.5 ±0.8 nM and all variant proteins displayed Kd values within ∼2-fold of this value (**Figure 2C**). Interestingly, this assay suggested that the ΔHSM and ΔOB-ΔHSM variants bound DNA slightly more *tightly* than wild type (Kd = 1.2 ±0.4 nM and 1.2 ±0.3 nM respectively), whereas the ΔOB mutant binding was slightly weaker (Kd = 5.7 ±1.2 nM).

We then tested the ability of ssDNA to stimulate the ATPase activity of HELB using a coupled assay (see Methods for details). The basal ATPase activity of wild type HELB is very low but strongly stimulated by ssDNA (**Figure 2D**). The relationship between [ssDNA] and ATPase activity is hyperbolic with a maximal turnover value (k_cat_) of 16.4 ±0.9 sec^-1^ and the concentration of ssDNA required for half-maximal stimulation of ATPase (K_DNA_) is 20.1 ±3.8 µM (in nucleotides). The k_cat_ values for all three variant HELB constructs were nearly identical to wild type (within the range 15.2-16.9 sec^-1^), whereas all K_DNA_ values were smaller, consistent with apparently tighter ssDNA binding (**Figure 2D**). For ΔHSM this reduction was modest (K_DNA_ = 13.9 ±2.7 µM) whereas for ΔOB and ΔHSM-ΔOB the reduction was 5-7 fold (K_DNA_ = 4.2 ±0.7 µM and 3.1 ±0.4 µM respectively).

In summary, these data show that all variant proteins retain the ability to bind ssDNA tightly, in some cases better than wild type, and to couple this binding to activation of ATP hydrolysis. Importantly, these data strongly suggest that all variant proteins are correctly folded as they retain the ssDNA-dependent ATPase activity associated with the core helicase domains; an activity which is expected to be coupled to ssDNA translocation.

### HELB variants retain ssDNA translocase activity, but the OB domain is required for ssDNA looping

We next used an optical trap coupled to a confocal fluorescence microscope (C-Trap instrument) to monitor ATP-dependent DNA translocation and ssDNA looping. A 22.2 knt ssDNA molecule was first immobilized between two optically trapped beads and then incubated with quantum dot labelled HELB in the presence of ATP (see **Figure S3** for a schematic of the experimental set-up). As expected, wild type HELB bound to the ssDNA and, in the presence of ATP, translocated in a unidirectional fashion with an average rate and processivity of 74 ± 35 nt sec^-1^ (mean ± SD) and 9 ± 4 knt respectively (**Figure 3**). The translocation polarity cannot be determined from this experiment, but independent bulk experiments have shown previously that HELB moves in the 5′>3′ direction [21]. A striking feature of the wild type data is the presence of frequent decreases in the apparent end-to-end distance between the beads (**Figure 3A**, lower panels; **Figure S3C**). In previous work we interpreted these events as the formation of ssDNA loops which were also demonstrated using magnetic tweezers [21]. Because HELB is monomeric in free solution [21], we proposed that its ability to form such loops arises from having two distinct DNA binding sites: one that maintains a firm grip on the DNA while another is translocating, resulting in the ‘reeling-in’ of the ssDNA rather than a simple forward movement. All of the HELB variant proteins retained the ability to translocate efficiently on ssDNA with rates and processivities that were similar to wild type (**Figures 3B and 3C**). Thus, ΔOB translocates at an average rate of 75 ± 29 nt sec^-1^ with a processivity of 7 ± 5 knt, values very similar to those of wild type HELB. ΔHSM translocates at 102 ± 23 nt sec^-1^, which is 36% faster than HELB, and exhibits a higher processivity (12 ± 5 knt). ΔOB-ΔHSM also translocates slightly faster than wild type HELB (94 ± 32 nt sec^-1^) and shows long trajectories indistinguishable in length from those of ΔOB (7 ± 5 knt). The most striking result however, is that the ssDNA looping events are almost completely absent from the traces for the ΔOB and ΔOB-ΔHSM proteins, whereas the ΔHSM variant forms loops slightly more frequently than wild type and with similar sizes and rates (**Figure 3A** and **Figure S3D-F**). This is consistent with the idea that the OB domain acts as an anchor that, in combination with ATP-dependent DNA translocation, causes ssDNA looping in the wild-type enzyme. Supporting this model, the loop extrusion velocity is comparable to the translocation velocity of HELB, and the observed loop sizes fall well within the processivity range of the protein (**Figure S3E and S3F**).

**Figure 3:**
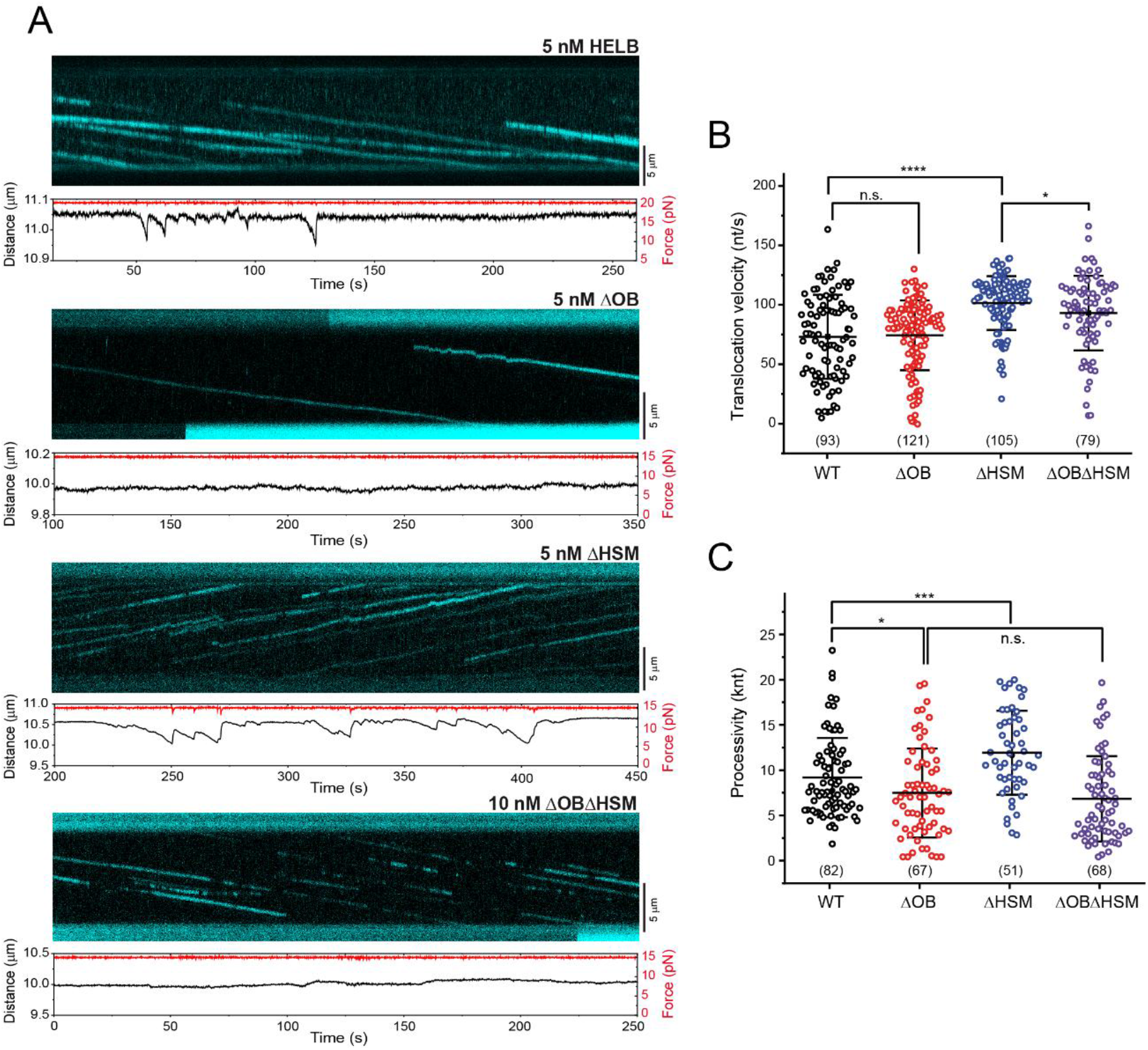
HELB variants retain ssDNA translocase activity, but the OB domain is required for ssDNA looping. (A) Sample kymographs showing translocation of quantum dot labelled HELB or HELB variants on ssDNA. The experiments were performed under a force-clamped condition at 15 pN and the lower panel for each trace shows the force and bead end-to-end distance with time. Note that wild type traces are characterised by frequent decreases in the bead end-to-end distance that are absent in traces for variants lacking the OB domain. (B) and (C) Distributions of translocation velocity and processivity for HELB and HELB variants. The central bar represents the mean, and the error bars indicate the SD. Samples sizes are indicated below each dataset.

### The HELB variants show qualitatively different defects in their ability to remove RPA from ssDNA

We next used the C-Trap to visualize the action of wild type and HELB variants on single DNA molecules coated with fluorescent RPA. A 22.2 knt ssDNA molecule was immobilized, incubated in a channel containing fluorescent RPA^MB543^ (**Figure S4**), then transferred to a channel with HELB variants and ATP but no free RPA. We recorded kymographs along the DNA axis and RPA clearance activity was visualized as the appearance of dark, shaded regions along the DNA. From these data we measure an overall rate of RPA clearance and infer the translocation velocity of individual HELB motors by monitoring the leading edge of individual clearance events.

As demonstrated previously [21], wild type HELB (used at 10 nM) binds to the RPA-coated ssDNA and translocates along the ssDNA, displacing RPA in the process (**Figure 4A**). We do not have the resolution to determine whether the initial association of HELB is to ssDNA gaps between RPA molecules or directly to the nucleoprotein filament. The RPA clearance activity of HELB exhibits high processivity, with single trajectories covering thousands of nucleotides, such that a small number of HELB molecules can efficiently remove all the RPA within just a few minutes. The overall RPA fluorescence intensity decays exponentially with an observed rate k_off_ = 0.646 ± 0.008 min^-1^ (k_off_ ± SE) (N = 13 kymographs) (**Figure 4B and 4C**). This is much faster than the rate at which RPA unbinds passively and/or photobleaches from ssDNA (0.109 ± 0.002 min^-1^, N = 21) or the rate at which HELB removes yeast RPA (with which it does not interact, 0.326 ± 0.003 min^-1^, N = 19). The apparent translocation velocity of wild type HELB is 86 ± 20 nt s^-1^ (N = 123 trajectories) (**Figure 4D**) which matches the rate of translocation in the absence of RPA. The onset times of individual RPA clearance activities are exponentially distributed which also allows us to define an observed initiation rate (k_on_) for wild type HELB of 0.62 ± 0.15 min^-1^ (**Figure 4E**). Overall, we observe that wild type HELB clears a densely coated RPA-ssDNA nucleoprotein filament at the same rate as it translocates over bare ssDNA, indicating that human RPA does not pose any obstacle for the enzyme. Alongside the fluorescent signal, we monitored changes in the distance between the beads that arise as a result of ATP-dependent protein activity (**Figure 4A**, lower traces). We observe gradual decreases in distance, followed by a sudden recovery of the initial extension which is attributed to ssDNA loop formation as discussed above [21]. The rate of loop formation (89 ± 27 nt s^-1^ (N = 68)) was similar to the translocation velocity (**SFigure 4**) and the loops formed by HELB were large, often encompassing hundreds of nucleotides (**SFigure 4**).

**Figure 4:**
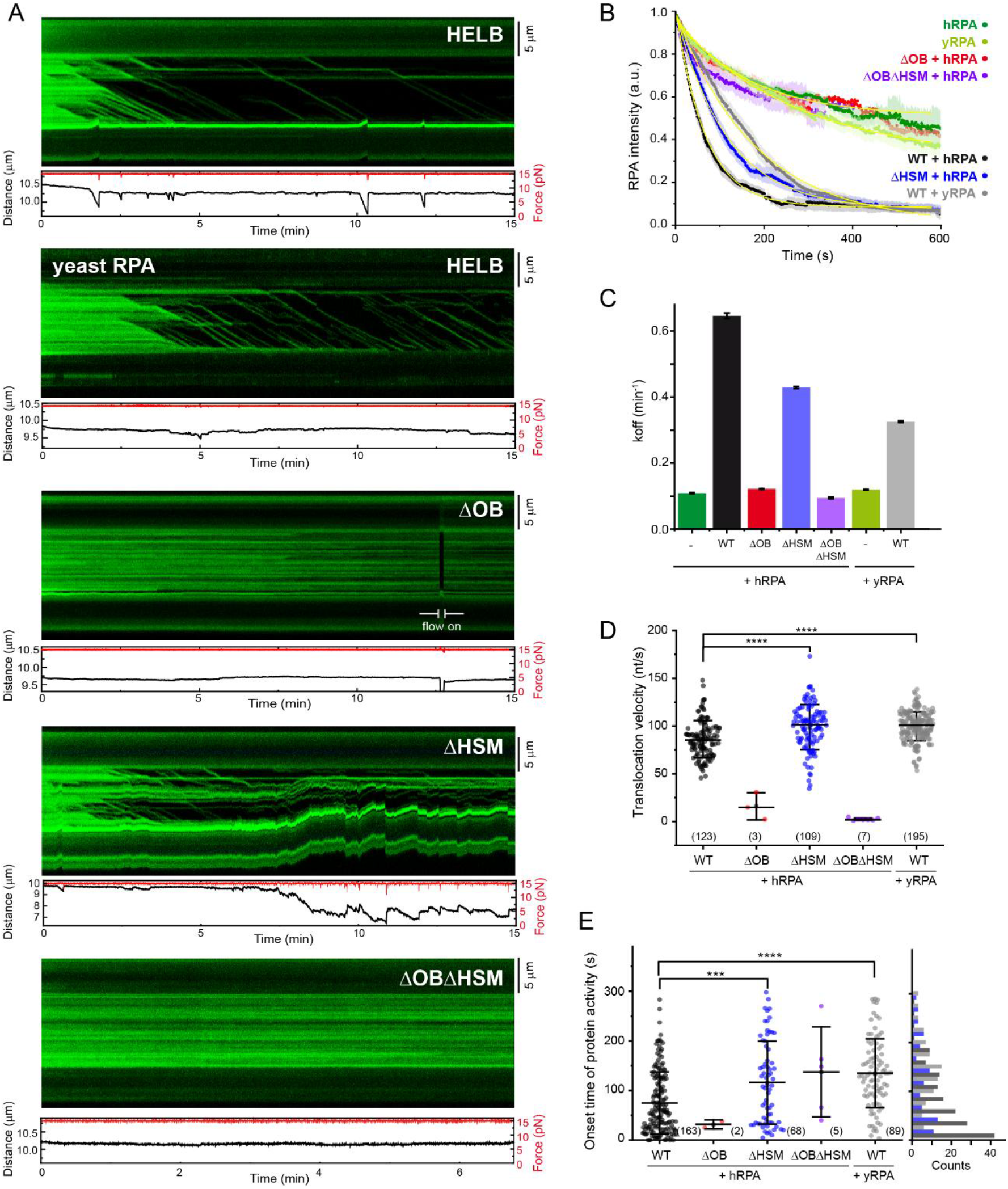
The HELB variants show qualitatively different defects in their ability to remove RPA from ssDNA. (A) Kymographs showing removal of fluorescently labelled RPA by 10 nM HELB and HELB variants. The lower panel shows the applied force and bead end-to-end distance. (B) Observed RPA fluorescence decrease induced by HELB and HELB variants. Traces are colour-coded according to the key and represent the mean decay over time in the total fluorescence intensity (i.e. the combined signal from all bound RPA molecules) measured along individual N DNA molecules or kymographs acquired under identical conditions (N indicated in the main text). Shaded regions represent the standard error of the mean. Yellow lines indicate exponential fits to the mean traces. Fits for variants lacking the OB fold are omitted for clarity to avoid overcrowding the plot. (C) Quantification of the observed rate of RPA clearance from the data shown in panel B. The k_off_ values are obtained from the exponential fits of average RPA decays at each condition. Error bars are the standard error of the fitting. (D) Implied translocation rate of HELB during RPA clearance. Individual data points are derived by monitoring the leading edge of RPA clearance events such as those shown in panel A. (E) Distribution of the onset time of protein activity for HELB and HELB variants. Each data point represents the time at which a processive RPA clearance event starts. For the interval scatter plots the centre bar represents the mean of the data and the error bars represent the SD. Sample size is indicated below the data; note that these are small for the ΔOB and ΔOBΔHSM variants due to the small number of productive RPA clearance events observed.

We next compared the activity of the ΔOB variant in this assay. At the same concentration as wild type (10 nM) RPA clearance events were rare and displayed very limited processivity (**Figure 4A**). The overall rate of RPA clearance was low (0.122 ± 0.002 min^-1^, N = 3), approaching the background value obtained for passive removal of RPA by simple unbinding and photobleaching (**Figure 4B**). The translocation velocity derived from the infrequent ΔOB trajectories was significantly lower than for the wild type, 16 ± 14 nt s^-1^ (N = 3) (**Figure 4D**) as was its processivity (**SFigure 4F**). These results show that deletion of the predicted OB-fold severely reduces the ability of the protein to catalyse RPA clearance, even though it can interact with, and translocate along, naked ssDNA normally. Importantly, and consistent with observations above, no loop formation was detected in the ΔOB experiments even under conditions where weak RPA clearance activity was observed (**Figure 4A**, lower panels). The inability of this HELB variant to form loops supports our earlier hypothesis that HELB possesses two DNA-binding sites, one of which is dependent upon the presence of the OB-fold.

We next measured RPA removal by the ΔHSM mutant at the same concentration (10 nM). We observed that ΔHSM binds and moves along ssDNA, following linear trajectories and clearing RPA in a broadly similar manner to wild type (**Figure 4A**). The RPA fluorescence decayed at a slower overall rate compared to wild type, but the effect was much less significant than with the OB mutant, yielding k_off_ = 0.429 ± 0.003 min^-1^ (N = 13) (**Figure 4**). Interestingly, analysis of the leading edge of individual RPA clearance events in kymographs showed that ΔHSM actually translocates on RPA filaments slightly *faster* than the wild type in the presence of RPA (99 ± 24 nt s^-1^ (N = 109)). Moreover, ΔHSM translocates long distances of 5 ± 3 knt while removing RPA. Importantly however, analysis of the onset times showed that ΔHSM activity was significantly delayed relative to wild type and no longer exponentially distributed (**Figure 4E**). This implies that successful initiation of RPA clearance requires multiple steps with comparable rate constants (for example, if multiple rounds of non-productive binding or abortive initiation were required before productive activity). Interestingly, the displacement of human RPA catalysed by the ΔHSM variant is broadly similar to displacement of yeast RPA by wild type HELB (translocation velocity 100 ± 15 nt s^-1^ (N = 195), processivity 12 ± 6 knt (N = 132) and intensity decay 0.326 ± 0.003 min^-1^) (**Figure 4 and SFigure 4).** Again, the reduced overall decay rate is not due to impaired displacement, but rather to a delayed onset of activity (compare ΔHSM/hRPA and WT/yRPA data in Figure 4E). In both cases therefore, it seems likely that the delayed RPA clearance observed reflects a failure to physically interact with the RPA complex. Consistent with the ssDNA translocation assays, we observe that the ΔHSM variant retains the ability to form DNA loops, as indicated by the distinctive changes in bead extension observed during RPA clearance (**Figure 4A**). These loops are formed at a similar rate and have a similar size to wild type (**SFigure 4**). In summary, our data show that the ΔHSM variant is defective for RPA clearance because of a slow initiation rate, but that once engaged with substrate, removal of RPA is similar or even better than wild type, leading overall to a modest decrease in the RPA clearance activity seen at this concentration of enzyme.

Finally, we monitored the RPA clearance activity of the double variant protein ΔOB- ΔHSM. RPA clearance events were infrequent, slow and exhibited minimal processivity (**Figures 4A-4D**). The overall clearance was very low (0.094 ± 0.003 min^-1^, N = 4), falling within the range of passive fluorescence decay observed in control experiments lacking HELB proteins. The overall activity of ΔOB-ΔHSM was so weak that the onset of activity could not be well defined due to a lack of data (**Figure 4E**). These results indicate that, although the motor activity of the double mutant remains intact and its ability to move on ssDNA is not impaired, the combined absence of the OB and HSM domains effectively render the protein inactive on RPA-ssDNA.

### Functional and physical interactions between HELB variants and RPA

To dissect the mechanistic basis for the impaired RPA clearance activities of the variant proteins we studied the direct binding of HELB to RPA alone or to RPA-ssDNA filaments, as well as the efficiency with which RPA-ssDNA filaments activated HELB ATPase activity.

To assess physical interaction between HELB and RPA in the absence of nucleic acids we used pulldown assays. StrepII-tagged HELB or HELB variants were used to bait Streptactin XT magnetic beads which were then tested for their ability to bind purified human RPA. Wild type HELB was able to pull down all three RPA subunits (**Figures 5A and 5B**). This direct protein-protein interaction is evidently weak, as it is not detectable by less sensitive approaches such as gel filtration (data not shown). The ΔOB construct retained the ability to pulldown RPA, albeit slightly impaired compared to wild type. In contrast, removal of either the HSM motif or of both the OB fold and the HSM motif resulted in a substantial, but incomplete, loss of the RPA interaction. These data show that the HSM motif is necessary for efficient interaction with RPA in the absence of ssDNA, but do not exclude the possibility that other regions of the protein also contribute.

**Figure 5:**
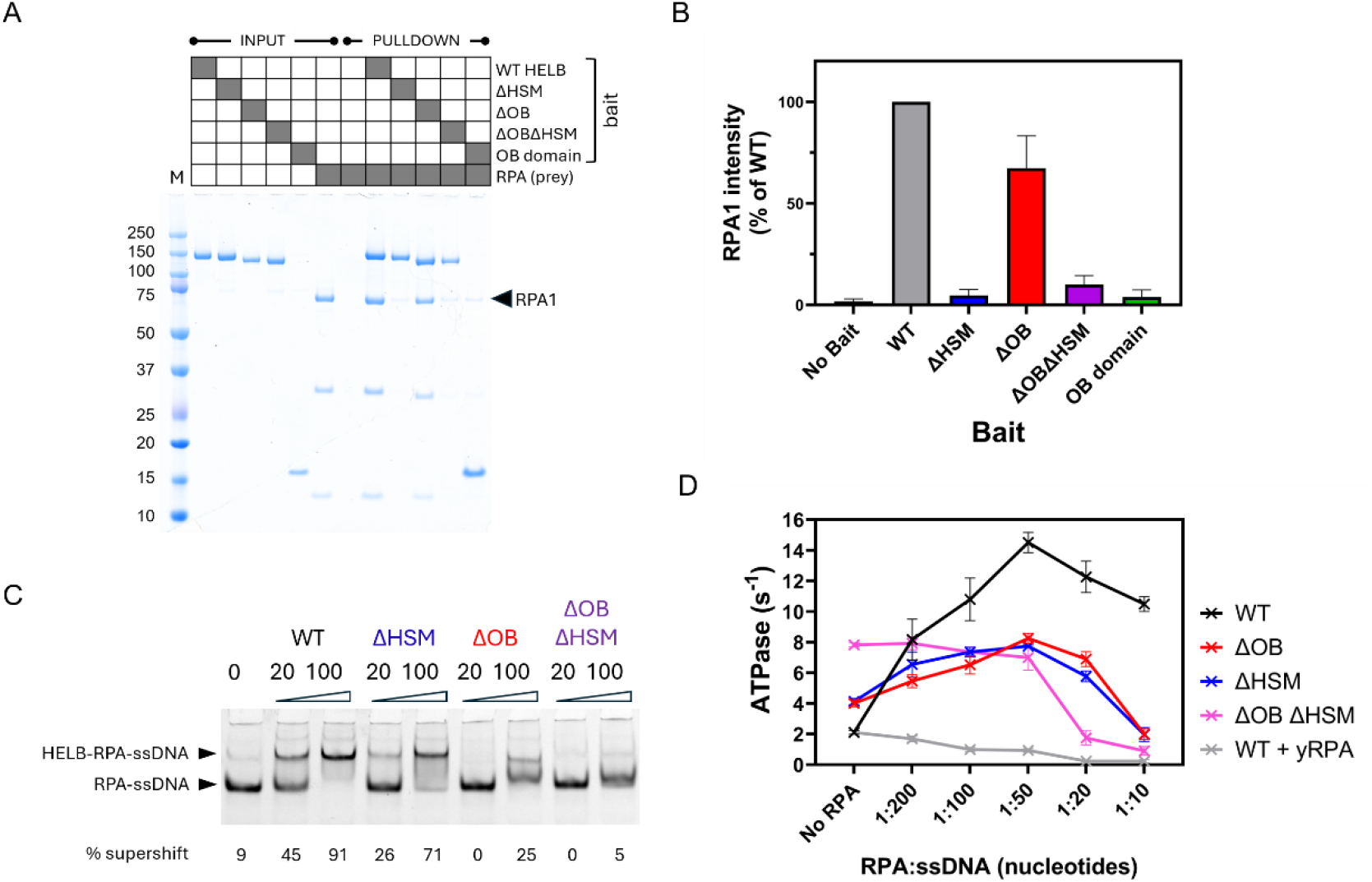
Functional and physical interactions between HELB variants and RPA. (A) RPA-pulldown assays using wild type or variant HELB proteins as baits in the absence of ssDNA. Pulldowns exploited the StrepII tag on the bait proteins. Loss of the HSM domain, but not of the OB domain, severely reduces pulldown of RPA from free solution. (B) Quantitation of the data shown in panel A. The signal intensity in the RPA1 band was normalised as a percentage of the wild type HELB-RPA1 pulldown. The data shown are the mean and standard deviation for three independent repeats. (C) RPA-ssDNA supershift assays were used to detect the formation of HELB-RPA-ssDNA ternary complexes. For the representative gel shown, the percentage of supershifted material is shown to provide a quantitative measure of the efficacy of HELB binding to RPA-ssDNA. A further example of this assay is shown in Figure S2 to demonstrate reproducibility. (D) Stimulation of ATPase activity by RPA for wild type and variant HELB proteins. These experiments are performed under conditions of low [ssDNA] (< K_DNA_) where addition of human RPA causes a strong ATPase stimulation for wild type HELB. Note that ssDNA is slightly over-saturated with RPA at 1 RPA heterotrimer per 10 nucleotides (see Methods for details) and that the x-axis scale is non-linear with the connecting line intended only as a guide for the eye.

We next performed EMSA supershift assays in which pre-formed complexes of RPA-ssDNA were titrated with wild type or variant HELB. The wild type HELB supershifts the RPA-ssDNA complex in a dose-dependent manner with most of labelled ssDNA appearing in a slowly migrating band at a concentration of 100 nM HELB (**Figures 5C and S Figure 2C**). We showed in previous studies [21] that these supershifted complexes contain HELB, RPA and ssDNA and do not form if human RPA is replaced with yeast RPA, strongly suggesting that we are measuring a specific recruitment event as opposed to the independent binding of HELB and RPA to a stretch of ssDNA. Deletion of the HSM motif moderately impairs binding of HELB to the filament. Deletion of the OB domain reduces the supershift to a greater extent, such that the majority of the RPA-ssDNA complex remaining unshifted at the highest concentration tested. Deletion of both the OB domain and the HSM motif nearly eliminates the formation of the supershifted band. Together the data show that both the HSM motif and the OB fold both contribute to stable binding of RPA-ssDNA filaments by HELB in the absence of ATP. This is in contrast with the pulldown experiments above, performed in the absence of DNA, where only the HSM motif appeared to play an important role in direct HELB-RPA protein-protein interaction. This implies that the full interaction between HELB and RPA filaments involves the remodelling of RPA into its ssDNA bound conformation.

Recruitment of HELB to RPA-ssDNA filaments can also be observed by monitoring the stimulation of ATPase activity afforded by the presence of RPA on the ssDNA substrate. As with all SF1 helicases, HELB ATPase activity is strongly stimulated by ssDNA due to tight coupling between ATP hydrolysis and DNA translocation (**Figure 2D**) [22]. At sub-saturating concentrations of ssDNA, where the observed rate of ATP hydrolysis is limited by the association rate of HELB with ssDNA, it is possible to observe a strong stimulatory effect of human RPA on ATPase activity, presumably because of its ability to recruit HELB to the ssDNA (**Figure 5D**, black trace). The ATPase activity increases with increasing [RPA] until a maximum stimulation of 7-fold is observed at a stoichiometry of 1 RPA heterotrimer per 50 ntds. However, a 5-fold stimulatory effect is still evident at saturating RPA (1 RPA per 10 ntds). This behaviour contrasts starkly with the effect of yeast RPA, which is inhibitory at all concentrations tested and eliminates ATPase activity at saturation (grey trace). The potent inhibitory effect of yeast RPA observed in bulk is not fully consistent with our observations of yeast RPA removal from single DNA molecules (**Figure 4**). We presume that, because RPA-filaments are held stretched out in a force clamped configuration in the C-trap instrument, and/or because the filament is transferred to an RPA-free buffer to perform the experiment, there may be small regions of accessible ssDNA that would not be present in bulk under zero applied force. Therefore, the single molecule RPA clearance assay may marginally reduce the need for HELB loading and underestimate the importance of HELB-RPA physical interactions for activity.

The OB and HSM variants (red and blue traces respectively) behave similarly to each other in this assay; we still observe stimulation of their ATPase activity by RPA but the effect is modest compared to wild type (at best ∼2-fold at 1 RPA per 50 ntds) and at saturation the RPA becomes inhibitory. Interestingly, removal of both the OB and the HSM domains (magenta trace) completely eliminates any stimulatory effect of RPA on the system. As was the case with wild type HELB and yeast RPA (grey trace), titration of the ssDNA binding protein results in a dose-dependent inhibition of ATPase presumably as the result of simple competition for the ssDNA. Overall, these data re-enforce our earlier observation (**Figure 2D**) that ssDNA-dependent ATPase is better than wild-type for all variants (particularly the double deletion variant) under conditions where RPA is absent and [ssDNA] is limiting. They show that the OB and HSM domains prevent access of the core motor domains to naked ssDNA yet facilitate loading and activation on saturated RPA filaments, consistent with a regulatory role for these structural elements.

## DISCUSSION

In this study, we explored the role of key structural elements of HELB in clearing RPA from ssDNA focusing on two structural regions, a predicted OB-fold and the HELB-specific motif. We showed that removal of either (or both) of these elements prevented efficient removal of RPA from ssDNA. The RPA clearance activity of HELB is entirely dependent on the helicase motor domains because inactivation of this region of the protein by mutation of the Walker A motif (K481A) eliminates all observed activity [21]. However, the RPA removal defects we observed in the OB and HSM variants cannot be explained by a lack of DNA motor activity as they retain ssDNA binding, ssDNA-dependent ATPase and translocase activities at levels equivalent to, or better than, wild type. Instead, these variants are specifically defective in their interactions with RPA and/or RPA-filaments. The HSM variant was unable to detectably interact with RPA in free solution and formed weak interactions with RPA-ssDNA filaments (as measured either directly or via RPA-ssDNA-dependent ATPase). Importantly, direct observation of RPA clearance activity showed that slower RPA removal resulted primarily from a slow initiation rate and that, once engaged with ssDNA, HSM translocation was coupled to processive RPA clearance as effectively as wild type. These observations are consistent with the idea that the HELB-specific motif is important for the recruitment of the helicase to RPA-ssDNA filaments and are in broad agreement with data from several other studies [6, 25, 26].

The behaviour of the ΔOB variant was different. It retained the ability to interact directly with free RPA in solution but formed weak interactions with RPA-ssDNA filaments. Direct observation of RPA filaments revealed a very low RPA clearance activity overall and when OB-dependent activity was observed the rate and processivity were significantly reduced relative to wild type. Moreover, ssDNA looping events, which are observed in both the wild type and ΔHSM variant experiments, were almost completely absent from ΔOB traces in both RPA clearance and ssDNA translocation experiments. These data are consistent with the idea that the OB domain plays a direct role in the RPA clearance activity, and with the possibility that ssDNA looping may be important for effective RPA removal. Given that one well-established role for OB domains is ssDNA binding, a simple explanation for these data is that the HELB-OB forms a secondary DNA binding site which acts as an anchor during translocation by the core DNA motor domains. The OB fold would thereby also contribute to the overall stability of HELB on the RPA-ssDNA filament. The OB-fold may, through direct or indirect interactions with RPA or nearby DNA, induce conformational or positional changes that reduce RPA stability on ssDNA and facilitate its displacement. It is important to note, however, that the apparent DNA binding activity we observe for purified HELB-OB domain alone is weak, and we cannot exclude the possibility that it is caused by a tight DNA binding contaminant. Therefore, it is important to consider alternative explanations for the loss of looping caused by deletion of the putative OB domain, such as an allosteric effect which triggers loss of DNA anchoring at an unknown distal site. Moreover, and although we have no direct evidence to support the idea, we cannot completely exclude the possibility that looping results from HELB oligomerisation or collision, and that this is OB-dependent. A structure of the HELB-RPA-ssDNA complex will be important to resolve these questions.

A speculative ‘working model’ for HELB recruitment and activation on RPA-ssDNA filaments based on our data is shown in **Figure 6**. The core motor domains (yellow arrow) of HELB are initially unable to engage ssDNA that is saturated with RPA (green). Therefore, initial binding of HELB to the exterior of the saturated RPA filament is mediated through the HSM domain (blue star) interaction with RPA70N to form an ‘encounter’ complex (**Figure 6, step 1**). This would increase competition for ssDNA binding between DNA binding elements in HELB and RPA, by increasing the local concentration of HELB, thereby facilitating full engagement of the HELB core motor and OB (red circle) domains with ssDNA (**Figure 6, step 2**). Alternatively, or additionally, HELB-RPA interaction may help to weaken RPA-ssDNA binding to allow dynamic exchange with HELB as has been suggested for other systems [16]. Once bound, the core motor and OB anchor domain form a functional ssDNA looping and RPA displacement machine (**Figure 6, step 3**).

**Figure 6:**
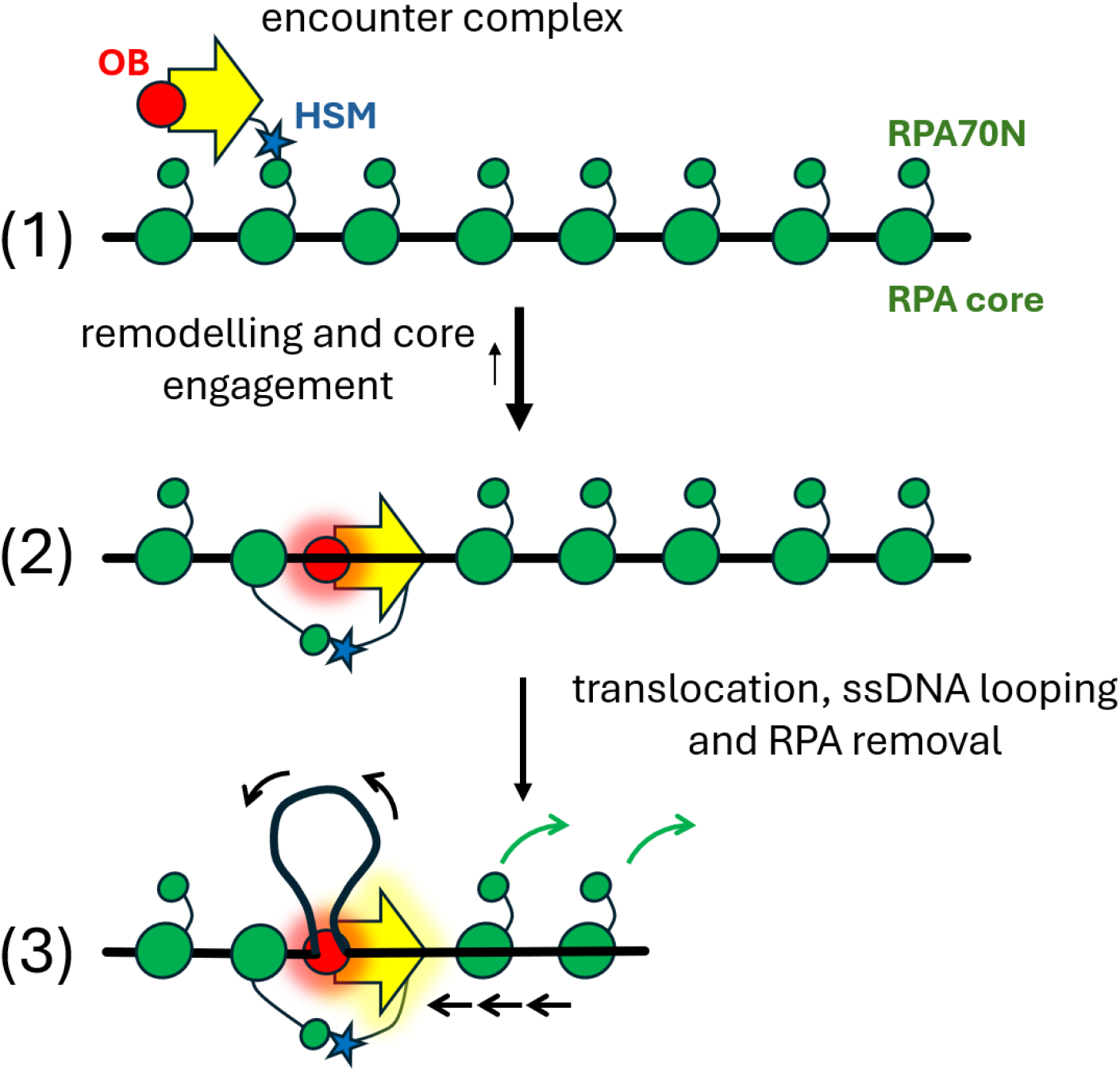
**A working model for HELB recruitment and activation on RPA-ssDNA filaments**. Step 1: Initial recruitment of HELB to the RPA filament mediated through HSM interaction with RPA70N forming an encounter complex. Step 2: Increased local HELB concentration promotes competition for ssDNA binding between RPA molecules and the DNA binding elements in HELB facilitating full engagement of the HELB core motor and OB domains with ssDNA. Step 3: The activated core motor domains and the OB anchor act as a ssDNA looping and RPA displacement machine. See text for further detail and discussion.

Removal of the OB and HSM domains results in inefficient RPA clearance but for different reasons (**Figure S5**), such that there is a synergistic detrimental effect when removing both. We suggest that lack of the HSM domain hinders the initial binding to the filament exterior, whereas loss of the OB domain both (a) prevents the remodelling that allows the core motor domains to stably engage with ssDNA and initiate translocation and (b) removes an anchor that facilitates ssDNA looping. It is possible that the remodelling event we hypothesise after initial engagement with the RPA filament involves interactions between the OB domain and RPA (**SFigure 1**), and/or competition for the underlying ssDNA between the OB domain and equivalent OB domains in RPA. These questions will be investigated in future studies. It is significant that we observed elevated ATPase activity of the ΔOB and ΔHSM variants on naked ssDNA substrates, implying that these structural features not only direct HELB to its correct substrate but also prevent erroneous activity on ssDNA lacking RPA. This regulation of HELB activity may help prevent unrestrained DNA translocation and/or unwinding activity that would otherwise be toxic. Indeed, loading and activation of SF1 DNA helicases at their intended substrate is a common feature of this enzyme class [22, 28].

HELB is one of several human helicases which have been shown to interact functionally and/or physically with RPA [15]. This close relationship is unsurprising, given that all *bona fide* helicases interact with single-stranded DNA and that RPA binds rapidly and tightly to exposed ssDNA in the cell. Among the best studied systems are the WRN, BLM and FANCJ helicases, all involved in DNA repair, where RPA is important for stimulation of DNA unwinding activity [29–34]. All three of these enzymes interact directly with RPA subunits and, in some cases, those physical interactions have been shown to be important for the stimulatory effect. We do not think these systems are directly analogous to HELB. Unlike the examples discussed above, we found that HELB is a relatively poor DNA unwinding enzyme and that this weak activity is inhibited rather than stimulated by RPA [21]. Instead, HELB seems to be recruited to RPA-ssDNA where its most prominent biochemical activity is to efficiently remove the RPA concomitant with translocation along the underlying ssDNA polymer. This property contrasts strikingly with the BLM helicase for example, where human RPA was shown to prevent BLM from binding or translocating on ssDNA using assays similar to those applied above [35]. The RPA clearance activity of HELB may facilitate the hand-off of ssDNA intermediates for further processing in a variety of DNA replication or repair processes. Alternatively (or additionally), given that RPA exhaustion has emerged as a driver of genomic instability and programmed cell death, HELB activity might also help to recycle RPA complexes between DNA damage at different loci [36, 37]. However, the links between HELB biochemical properties and its cellular function(s) remain poorly defined, not least because the precise roles of HELB in cells have been the subject of relatively few studies, some of which are contradictory.

This work has demonstrated the importance of the OB and HSM domains in HELB for recruitment and activation at RPA-ssDNA filaments. Many of the human SNPs associated with fertility traits map to these precise regions of the HELB structure (D117Y, D506G and E522D; [8, 11, 26]). Therefore, our data also suggest a mechanistic basis for the link between HELB mutations and their impact on human health, namely the failure to efficiently remove RPA from ssDNA.

## MATERIALS AND METHODS

### Purification of HELB variants

A synthetic gene codon-optimised for *S. frugiperda* encoding wild type human HELB (Geneart, Invitrogen) was cloned into the pACEBac1 vector using the BamHI and XbaI restriction sites for use in the MultiBac system (Geneva Biotech). Following cleavage of a construct with a C-terminal 3C-cleavable StrepII tag, we obtained HELB in good yield and purity. The full-length recombinant protein as prepared (i.e. post-3C cleavage) contains a C-terminal extension-SGLEVLFQ (MW = 124,126 Da monomer) with the rest of the protein identical to UniProt entry Q8NG08. Mutagenesis (Quikchange XL, Agilent) was performed using this construct to create the ΔOB, ΔHSM and ΔOB-ΔHSM truncation mutants. For the ΔOB construct, the DNA coding for amino acids 1-117 was removed from the vector. For the ΔHSM the DNA coding for amino acids 496-531 was replaced with a short linker coding for GGSGSG. For the ΔOB-ΔHSM construct both modifications were made in the same vector.

Bacmids were prepared by transposition of these plasmids and were used to transfect Sf9 cells in ESF921 (Expression Systems LLC) before viral amplification in the same cell line. For large scale expression, 500 mL of Sf9 or Hi5 cells at density 2 x 10^6^ /mL were infected with 25 mL of P3 virus and harvested by centrifugation after 70 hours at 27°C with shaking. The pellets were lysed into buffer containing 50 mM Tris pH 8.0, 100 mM NaCl, 1 mM TCEP, 10% glycerol, protease inhibitor cocktail (Roche) and then sonicated on ice for a total of 2 minutes. After centrifugation at 4°C for 30 mins at 50000g, the cleared lysate was applied to Streptactin-XT beads (GE Healthcare) in batch and incubated for 1 h at 4°C with rotation. After washing twice in batch with buffer containing 20 mM Tris HCl pH 8.0, 100 mM NaCl, 5% glycerol, 0.5 mM TCEP, once with the same buffer containing 1M NaCl, and twice again with buffer containing 20 mM Tris HCl pH 8.0, 100 mM NaCl, 5% glycerol, 0.5 mM TCEP, the protein was then eluted in the same buffer containing 50 mM biotin. The HELB-containing fractions were then applied to a 5 mL Heparin column (GE Healthcare). After washing, HELB was eluted with a gradient from 100 mM to 1 M NaCl over 16 CV in buffer containing 20 mM Tris HCl pH 8.0, 0.5 mM TCEP. The HELB-containing fractions were pooled and digested for 3h at 4°C with 3C protease to remove the StrepII tag (in some cases, to deliberately recover tagged protein for pulldown studies, this cleavage step was omitted). The cleavage reaction was run over a 5 mL Streptactin column (Qiagen) to remove any uncleaved HELB and free StrepII peptide and the cleaved HELB-containing flow-through collected. This material was spin concentrated (50,000 Da cutoff, Millipore) to a volume of approximately 200 µL and applied to a Superose6 10/300 column in buffer containing 20 mM Tris HCl pH 8.0, 200 mM NaCl, 0.5 mM TCEP. The HELB peak eluted after a volume of 15.5 mL, and was then concentrated using a centrifugal filter unit (Millipore) with glycerol added to a final concentration of 10% (v/v) before the protein was stored at-80°C. Protein concentration was determined using theoretical extinction coefficients of 120,780 M^-1^ cm^-1^ (wild type), 102,330 M^-1^ cm^-1^ (ΔOB), 115,280 M-1 cm-1 (ΔHSM), 96830 M-1 cm-1 (ΔOB ΔHSM). In the cases where the StrepII tag was left intact, the extinction coefficient increases by 5500 M^-1^ cm^-1^ in each case.

A construct encoding the HELB OB domain alone (amino acids 1-115) was tagged and expressed in the same way as the other variants but was prepared as follows: the pellets were lysed into buffer containing 50 mM Tris pH 8.0, 500 mM NaCl, 1 mM TCEP, 10% glycerol, protease inhibitor cocktail (Roche) and then sonicated on ice for a total of 2 minutes. After centrifugation at 4°C for 30 mins at 50000g, the cleared lysate was applied to Streptactin-XT beads (GE Healthcare) in batch and incubated for 1 h at 4°C with rotation. After washing five times in batch with buffer containing 20 mM Tris HCl pH 8.0, 500 mM NaCl, 5% glycerol, 0.5 mM TCEP, once with the same buffer containing 1M NaCl, and twice again with buffer containing 20 mM Tris HCl pH 8.0, 500 mM NaCl, 5% glycerol, 0.5 mM TCEP, the protein was then eluted in the same buffer containing 50 mM biotin. This was spin concentrated (10,000 Da cutoff, Millipore) to a volume of approximately 200 µL and applied to a Superose6 10/300 column in buffer containing 20 mM Tris HCl pH 8.0, 200 mM NaCl, 0.5 mM TCEP. The HELB OB peak eluted after a volume of 17 mL and was then concentrated using a centrifugal filter unit (Millipore) with glycerol added to a final concentration of 10% (v/v) and the protein stored at-80°C. Protein concentration was determined using a theoretical extinction coefficient of 23,950 _M-1 cm-1._

To produce biotinylated HELB and variant proteins, a modified version of the wild type HELB pACEBac1 plasmid was constructed containing a C-terminal Avi-tag upstream of the cleavable StrepII tag. The proteins were expressed and purified in an identical manner to wild type. Following StrepII-tag cleavage, the HELB-Avi protein was treated with BirA and biotin to site-specifically biotinylate the protein according to a published protocol [38] with the following minor modifications. A reaction of 50 µM HELB, 2 µM BirA, 1 mM biotin, 5 mM MgCl_2_ and 5 mM ATP was incubated at 4°C for 1 hour before being injected onto a Superose 6 10/300 column to further purify HELB and remove the other reagents. After SEC purification according to the wild type method, the protein was concentrated and biotinylation efficiency was estimated at ∼100% by PAGE gel. Conjugation of biotinylated HELB protein to streptavidin did not affect its biochemical activities [21]. In the cases where the StrepII tag was not removed, the pooled heparin fractions were concentrated and injected straight onto a Superose6 SEC column.

### Electrophoretic mobility shift and supershift assays

5′-Cy5-labelled single-stranded 25mer DNA (5 nM final) was mixed with increasing amounts of HELB protein in a total volume of 10 μL 1X EMSA buffer (20 mM Tris HCl pH 8.0, 100 mM NaCl, 1 mM DTT, 0.1 mg/mL BSA, 5% glycerol) and then incubated for 10 min at 25°C. In the cases where RPA was used, 20 nM final concentration was added to the DNA before incubation with HELB. The samples were then loaded onto a native 6% polyacrylamide (29:1 acrylamide:bis-acrylamide ratio) 1xTBE gel and separated by electrophoresis in 1xTBE at 150V for 35 minutes. The gels were visualised using a Typhoon scanner and analysed using the ROI tools in Fiji [39]. Quantitation of the supershift was performed as follows: after background subtraction, the signal intensity for each lane spanning the entire unshifted and supershifted DNA region was first measured for each condition. Subsequently, the intensity of the well-defined unshifted band was measured. Supershifted DNA is defined as the percentage of the total signal that is not present in this unshifted lower band.

### Protein-induced fluorescence enhancement (PIFE) DNA binding assays

3′-Cy3-labelled 40mer poly(dT) ssDNA substrate (2 nM final) was mixed in a quartz cuvette (Hellma) with increasing amounts of HELB protein in a total volume of 150 μL 1X PIFE buffer (20 mM Tris HCl pH 8.0, 100 mM NaCl, 1 mM DTT). After 1 minute, the PIFE signal is stable and the protein DNA mixture was scanned three times (λEx=530 nm, λEm=562 nm, 5 nm slit widths) allowing the average fluorescence intensity to be measured. The fully saturated DNA with wild type HELB gave an increase in fluorescence of around 60% compared to the original signal of free DNA. For each titration, saturation of the PIFE signal was achieved and the measurements were normalised with the initial reading being 0 and the highest reading being 100%. Data were fit using Prism (GraphPad) to the following equation for tight binding which assumes a stoichiometry of 1 HELB to each DNA molecule:

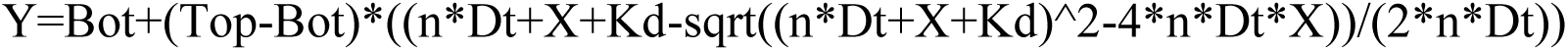

Where Y = normalized PIFE signal; X = total protein concentration (nM) added in the titration; Bot = signal with no protein bound. Top = signal at saturation of the DNA binding sites. Kd = dissociation constant. Dt = total DNA concentration (fixed at 2 nM); n = HELB binding sites per DNA molecule (fixed at 1).

### ATPase assays

ATPase activity was measured by coupling the hydrolysis of ATP to the oxidation of NADH which gives a change in absorbance at 340 nm. Reactions were performed in a buffer containing 20 mM Tris-HCl pH 8.0, 50 mM NaCl, 1 mM DTT, 5 mM MgCl_2_, 50 U/mL lactate dehydrogenase, 50 U/mL pyruvate dehydrogenase, 1 mM PEP and 100 µg/mL NADH. The concentration of HELB was 50 nM in these assays. Rates of ATP hydrolysis were measured over 1-5 min at 25°C. For calculation of K_DNA_, varying concentrations of ΦX174 virion ssDNA was used with the ATP concentration fixed at 4 mM. Data wre fit using a variation of the weak binding equation in Prism (GraphPad).

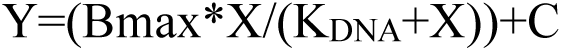

Where Y = observed ATPase rate, X = total ssDNA concentration added in the titration; Bmax = ATPase rate at saturating (ssDNA], K_DNA_ = apparent affinity for ssDNA; C = offset to allow for non-zero ATPase activity at zero ssDNA.

In the cases where RPA was used to assemble a nucleoprotein filament, the composition of the substrates was calculated based on the molar ratio of RPA-to-DNA. ΦX174 virion ssDNA was fixed at a final concentration of 2µM throughout the assays, with increasing amounts of RPA pre-incubated with the ssDNA before adding to the cuvette. For example, the 1 in 100 substrate was formed by mixing an equal volume of 100 µM ssDNA with 1 µM RPA to give 1 RPA heterotrimer per 100 nucleotides. 2uL of this was then added to the cuvette (100uL total volume) to give 2 µM of substrate as defined by the ssDNA. References to RPA saturation are based on the assumption that a single RPA heterotrimer binds to ssDNA with a footprint of approximately 20-30 nucleotides.

### RPA interaction pull-downs

To assess the interactions of the HELB variants with RPA we used a pull-down assay utilising the C-terminal StrepII tag on HELB. Each condition used 10 uL of Streptavidin-XT beads which were pre-equilibrated in binding buffer (25 mM Tris-Acetate pH7.5, 50 mM KCl, 1 mM TCEP, 10% glycerol, 0.1 mg/mL BSA, 0.01% Triton-X100). Beads were incubated with 2 μg HELB or HELB variants in 50μL binding buffer for 30 minutes at 4°C with mixing by rotation. After washing twice with 100uL of binding buffer, the beads were incubated with 2.5 μg RPA in 50 μL binding buffer for 30 minutes at 4°C with mixing by rotation. After washing twice with 100 uL of binding buffer with BSA omitted, beads were incubated with 15 uL 4X Laemmelli Buffer for 5 mins. This buffer was removed and boiled for 3 minutes at 95°C before loading onto an SDS-PAGE gel (Thermo Bolt Bis-Tris Plus) and running according to the manufacturer’s instructions. The gel was visualised using Coomassie blue. For quantitation, the ROI tools application in Fiji was used to determine the total intensity of the RPA1 band for each pulldown condition [39].

### DNA substrates for single-molecule experiments

To obtain long ssDNA tethers, we fabricated a 22.2 kbp dsDNA construct that incorporated nicks and a dephosphorylated handle in one of the DNA strands, following the detailed protocol described in [40]. Briefly, the central part of the DNA construct was obtained by digestion with XhoI and NotI (New England Biolabs) of a large homemade plasmid that contained three BbvCI restriction sites. The fragment was ligated to two highly biotinylated handles of ∼1 kbps ending in NotI or XhoI, being the NotI handle dephosphorylated. This creates a nick in one of the strands. Then, the dsDNA molecules were digested with the nicking enzyme Nb.BbvCI (New England Biolabs) leading to a final DNA construct with 4 nicks in the same strand, to facilitate the process of removal of this nicked strand by pulling in the optical tweezers setup.

### Optical tweezers combined with confocal microscopy

We used an instrument that integrates confocal fluorescence microscopy with dual-trap optical tweezers and microfluidics (C-Trap, LUMICKS). The experiments were conducted in a microfluidic cell with five independent channels (**SFigure 3 and SFigure 4**). Prior to the experiments, channels 4 and 5, which would contain proteins, were passivated by flowing bovine serum albumin (BSA, 1 mg/mL in PBS, Sigma-Aldrich) for 30 min. Streptavidin-coated polystyrene beads (0.005% w/v, 4.34 μm, Spherotech), 22.2-kbp-dsDNA (10 pM) molecules intended for ssDNA fabrication and melting buffer were introduced into channels 1-3, respectively. Melting buffer consisted on 10 mM Tris pH7.5, 25 mM KCl, 10 mM DTT, 1 mM CaCl_2_. The laminar flow within the channels prevents content mixing, allowing the optical traps to move throughout the channels and sequentially assemble the experimental configuration. A single DNA tether was caught between two streptavidin beads in the optical traps and moved to channel 3. Here, the duplex DNA was subjected to forces over the overstretching transition, leading to the removal of the nicked strand. The presence of a ssDNA was verified by force-extension curves. The ssDNA tether was moved to channel 4 and 5 for protein loading and imaging in activity buffer: 25 mM Tris pH 7.5, 100 mM KCl, 10 mM DTT, 5 mM MgCl_2_ and 5 mM CaCl_2_. For experiments to assess the activity in the absence of RPA, channel 4 contained 5 or 10 nM HELB or HELB variants conjugated with QDs in activity buffer supplemented with 2 mM ATP. For experiments on RPA-ssDNA, ssDNA tethers were first incubated inside channel 4 containing 15 nM RPA^MB543^ for 2 min (at F = 15 pN). The RPA-coated ssDNA was then moved to channel 5 containing 10 nM HELB or variants in the same buffer supplemented with 2 mM ATP, an oxygen scavenging system (2.5 mM 3,4-dihydroxybenzoic acid, 250 nM protocatechuate dioxygenase) and 1 mM Trolox. All experiments were performed at 15 pN and at room temperature.

The optical trap was calibrated to achieve a stiffness of 0.3-0.4 pN/nm. For bare ssDNA experiments, biotinylated HELB and its variants were conjugated to streptavidin-coated quantum dots (Qdot^TM^ 525, Invitrogen) by incubating a 1:3 protein-to-QD mixture in a small volume on ice for 30 min prior to dilution and introduction into the C-Trap. Confocal images were acquired using a 488 nm laser at 5% power, with fluorescence emission collected through a 512/25 nm bandpass (blue) filter. For experiments with unlabelled HELB or its variants on RPA-ssDNA, confocal images were acquired using a 532 nm laser (human RPA^MB543^ or yeast RPA^Cy3^ excitation) at 5% power, with fluorescence emission detected through a 585/75 nm bandpass (green) filter. Kymographs were generated via confocal line scanning along the DNA, with a pixel dwell time of approximately 0.1 ms/px.

Data generated by the C-Trap in HDF5 format were processed to extract fluorescence intensity profiles from individual RPA-coated ssDNA tethers. Custom-written Python scripts were used to export kymograph data as ASCII matrix files, which were subsequently imported into a custom LabVIEW program for further analysis. To determine the fluorescence intensity over time for each DNA molecule, intensity was averaged every 30 frames (∼1.5s) and photon counts were summed along the DNA axis at each averaged time point. Intensity profiles and curve fitting were generated using OriginPro software. Translocation velocities and processivities were determined from linear trajectories using Fiji [39]. Force and extension data were analyzed using custom-written Python scripts and OriginPro.

## ACKNOWLEDGEMENTS

Work in the M.S.D. laboratory was supported by the Medical Research Council (grant number MR/Y012070/1). Research in the F. M.-H. laboratory was funded by the Agencia Estatal de Investigación (AEI/10.13039/501100011033) and the Ministerio de Ciencia, Innovación y Universidades, and co-funded by the European Regional Development Fund (ERDF-EU) through grant PID2023-146255NB-I00. Additional funding to the F.M.-H. laboratory was provided by the Autonomous Region of Madrid through the Dirección General de Investigación e Innovación Tecnológica under the R&D Activities Program (grant TEC-2024/TEC-158; TecNanoBio-CM). Work in the E.A. laboratory was supported by the National Institutes of Health (grant numbers R35GM149320 and R01CA305375).

## AUTHOR CONTRIBUTIONS

E.A., F.M.-H., and M.S.D. acquired funding. O.J.W., S.H., F.M.-H., and M.S.D. designed the research; O.J.W., S.H., C.A.-R., and A.M. performed the research and analysed data; E.A., F.M.-H., and M.S.D. supervised the research; O.J.W., S.H., F.M.-H., and M.S.D. wrote the manuscript. All authors reviewed and edited the manuscript.

## SUPPLEMENTARY FIGURES

**Figure S1:**
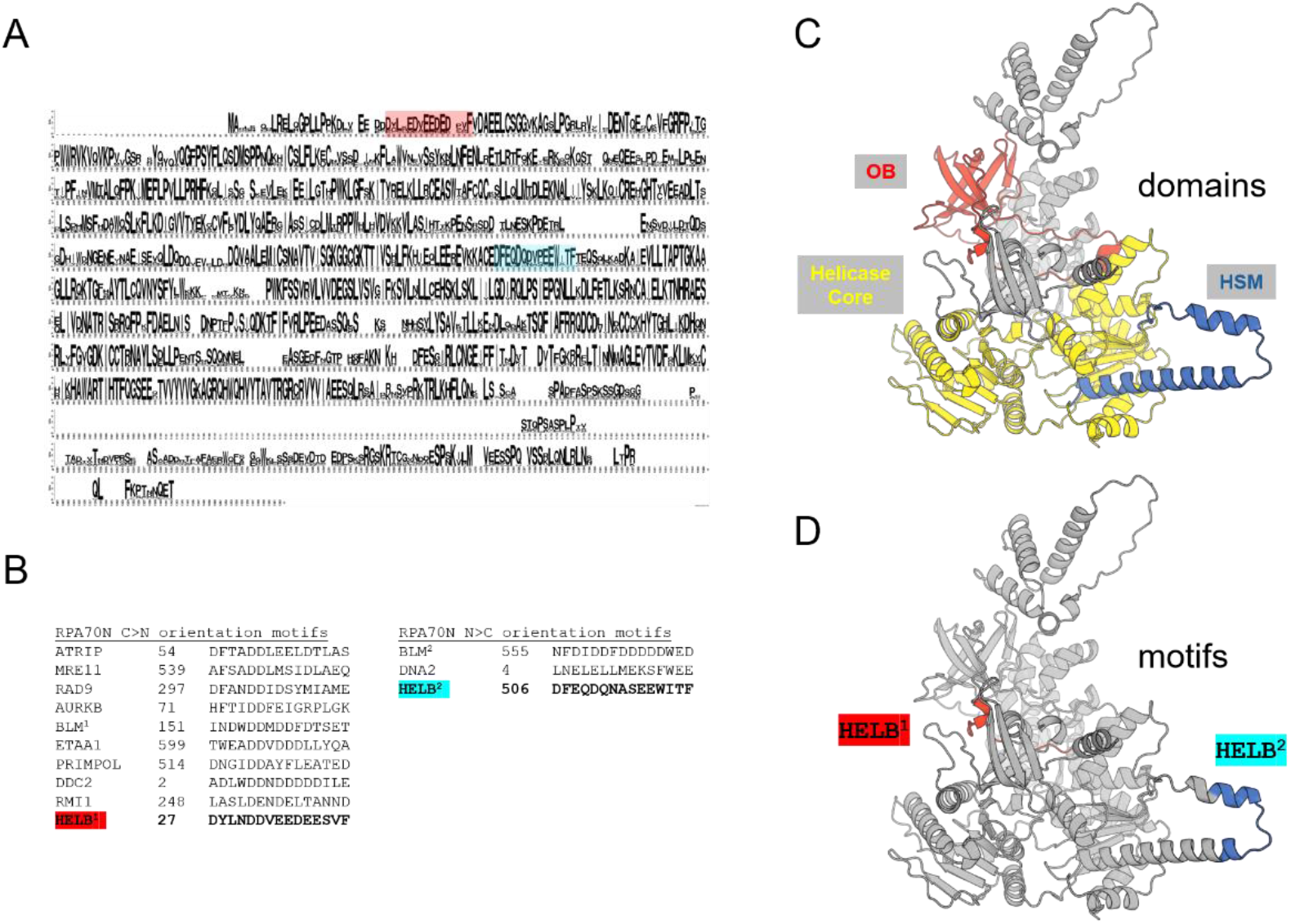
Putative RPA interaction motifs in HELB. (A) Possible RPA interaction motifs (highlighted red and blue) within a multiple sequence alignment of HELB homologues in WebLogo format [41]. To make the Weblogo, fifty representative HELB homologues were retrieved from the UniRef90 database using an E value cutoff of 1e^-5^ and then aligned in COBALT [42, 43]. (B) These conserved motifs contain many basic and hydrophobic residues which is characteristic of proteins that interact with the N-terminal region of RPA70. Note that binding of such peptides to the RPA70N domain has been shown to occur in diverse modes and in two orientations denoted here as N>C and C>N [25]. Therefore, the alignments between peptides are not necessarily intended to indicate conservation of function for individual residues. There is existing experimental support for the role of the HELB^2^ motif (blue) motif in RPA interaction [6, 25]. (C) AlphaFold3 model of HELB highlighting an OB-domain (red), the helicase core domains (yellow) and the HELB Specific Motif (HSM) (blue). (D) AlphaFold3 model of HELB highlighting positions of the putative RPA interaction motifs (colour-coding is as in panels A and B). The HELB^1^ motif is part of a short insert within the OB domain and the HELB^2^ motif resides ta the distal tip of the HSM domain.

**Figure S2:**
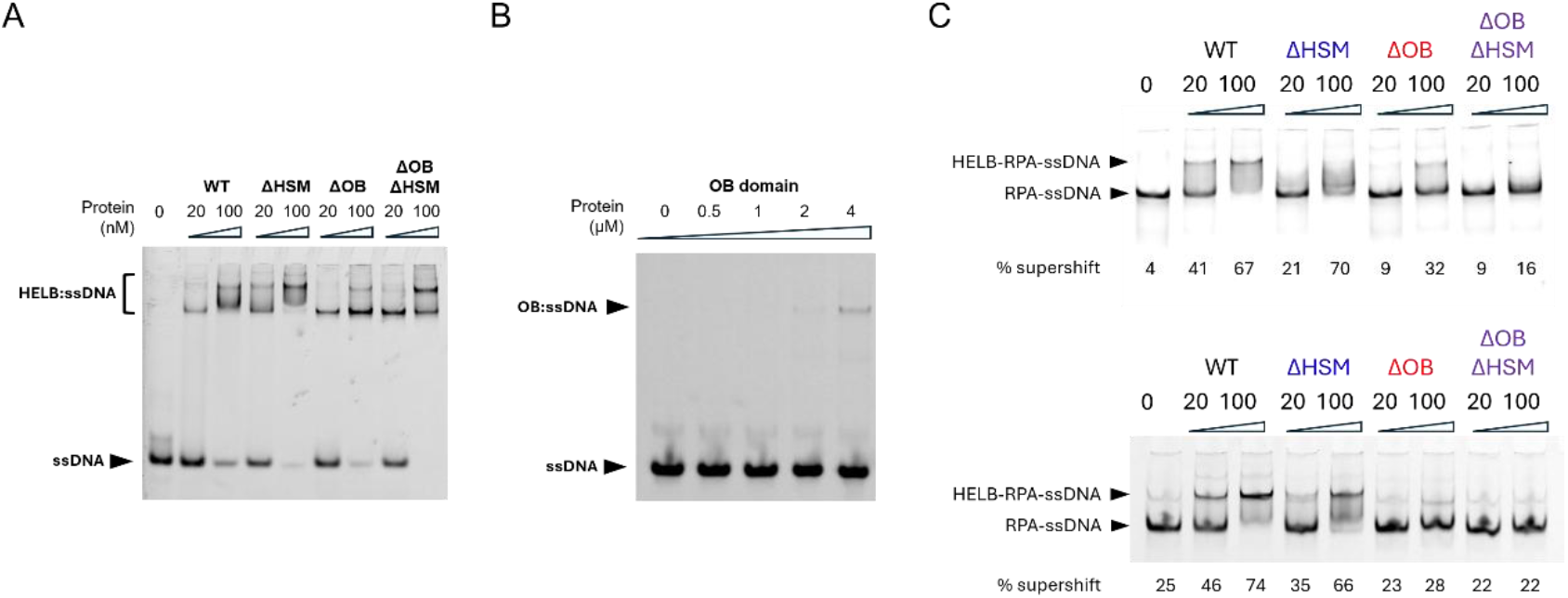
Further examples of ssDNA EMSA and RPA:ssDNA supershift assays for HELB variants. (A) Additional example of HELB EMSA experiment (equivalent to main paper Figure 2A). The gel shows binding of a 25mer ssDNA oligonucleotide (5 nM) by wild-type and variant HELB proteins at the concentrations indicated. (B) Additional example of HELB-OB EMSA experiment (equivalent to main paper Figure 2B). The gel shows binding of a 25mer ssDNA oligonucleotide (5 nM) by HELB-OB domain at the concentrations indicated. (C) Additional examples of HELB RPA:ssDNA complex supershifts (equivalent to main paper Figure 5C). The percentage of supershifted material is shown to provide a quantitative measure of the efficacy of HELB binding to RPA-ssDNA.

**Figure S3:**
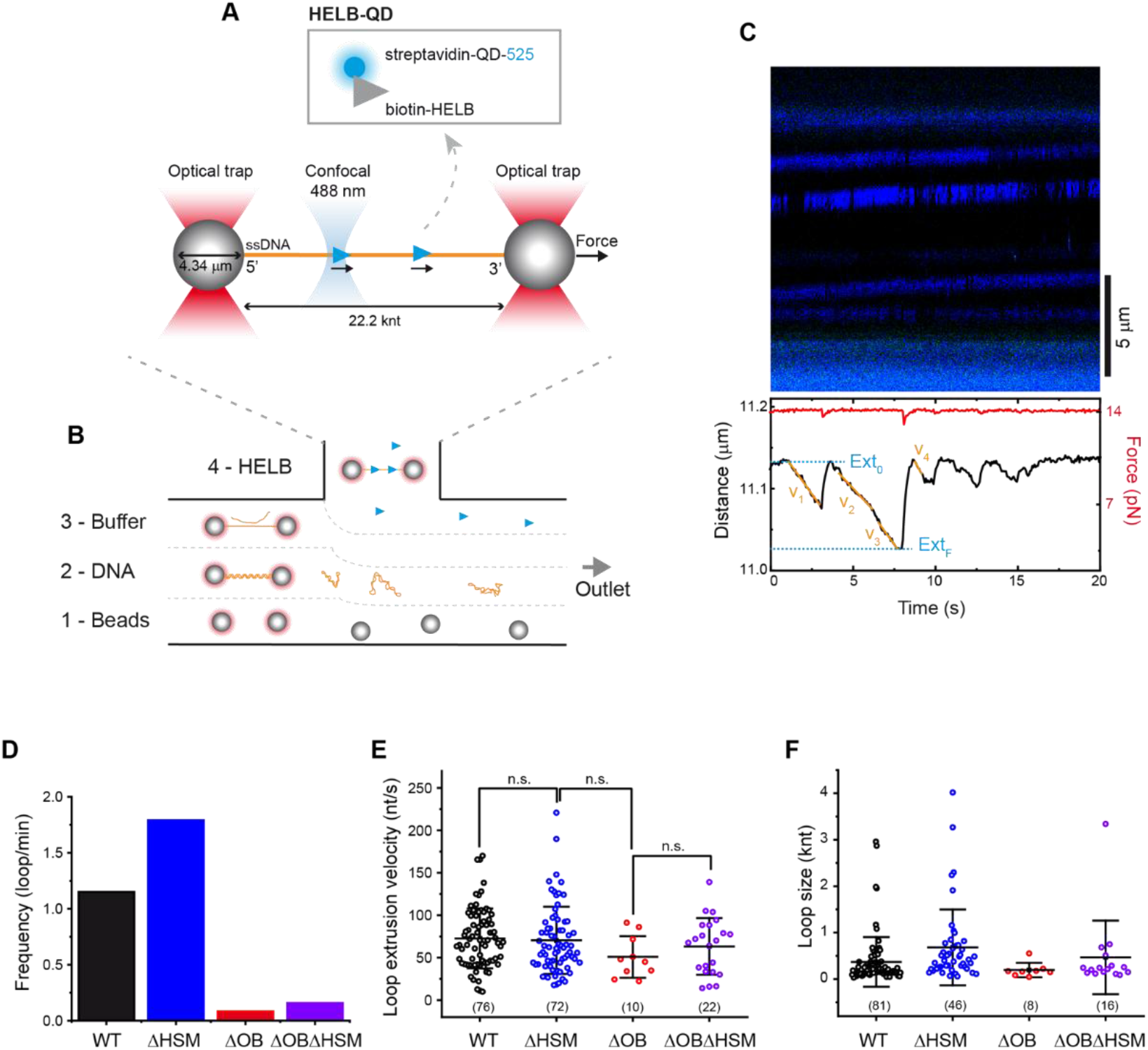
Schematic representation of the single-molecule DNA translocation assay for HELB. (A) Overview of the experimental configuration to measure the translocation of HELB and HELB variants using the C-Trap system. Single 22.2-knt ssDNA molecules were tethered between two streptavidin-coated beads held in dual optical traps. A confocal laser scanned the DNA tethers to visualize HELB molecules labelled with a quantum dot. (B) Schematic of the microfluidic flow cell employed in the optical tweezers measurements. Individual tethers were assembled in channels 1-3 under laminar flow conditions, with separate channels containing streptavidin-coated beads (1), DNA for ssDNA formation (2), and low-ionic-strength buffer (3). Once a ssDNA tether is formed, the traps were transferred to channel 4 for protein loading and fluorescent imaging. (C) Representative experiment showing HELB activity on ssDNA. Kymograph displaying translocating HELB trajectories (upper panel) and the corresponding DNA extension and force traces showing looping events (lower panel). The loop extrusion velocity was determined by linear regression of the segments corresponding to the gradual decrease in DNA extension (v_i_). Loop size was calculated as the difference between the initial extension and the minimum extension reached during a looping event (Ext_0_-Ext_F_). (D) Frequency of loop formation observed in the presence of HELB and HELB variants. (E) and (F) Loop extrusion velocity and loop size measured for 5 nM HELB and HELB variants on bare ssDNA. For the interval scatter plots, the central bar represents the mean and the error bars indicate the SD. Sample sizes are indicated below the plots.

**Figure S4:**
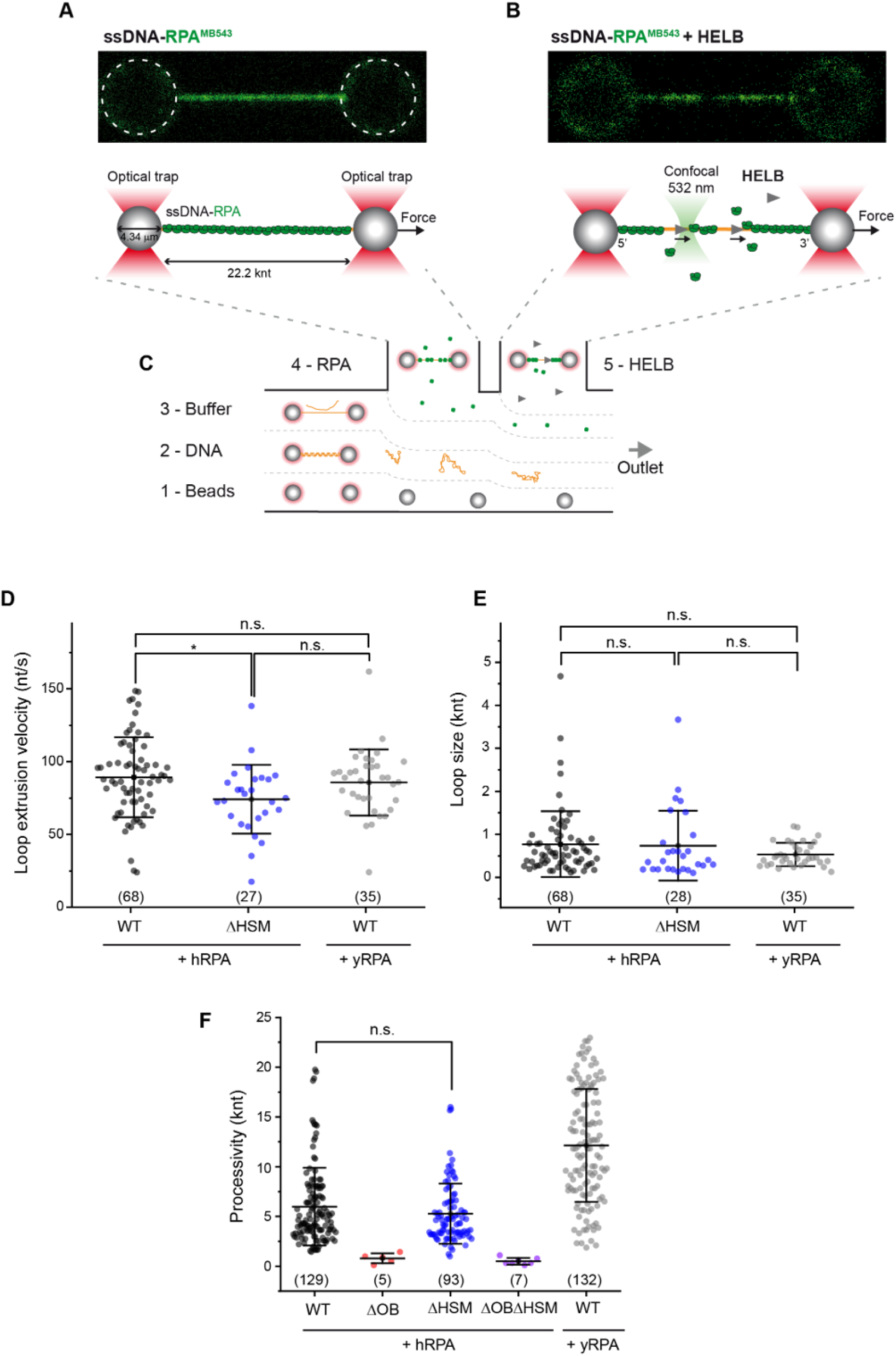
Schematic illustration of the single-molecule RPA clearance assay. (A) Confocal scan (upper panel) and schematic (lower panel) showing fluorescently labelled RPA bound to ssDNA tethered between two optically trapped beads in the C-Trap. The scan was acquired after 2 min in channel 4 containing 15 nM RPA-MB543. (B) The effect of HELB activity on RPA-ssDNA is monitor in channel 5 containing HELB and ATP but no RPA. (C) Schematic representation of the five-channel fluid cell used in C-Trap experiments. The optical traps are moved sequentially through channels containing a continuously maintained laminar flow according to the following workflow: (1) a pair of micrometre-sized beads is optically trapped in channel 1; (2) a dsDNA tether is immobilized in channel 2; (3) a ssDNA tether is generated by force-induced melting in channel 3; (4) the ssDNA is incubated with RPA in channel 4 containing 15 nM RPA-MB543 for 2 min; and (5) the activity of wild type HELB or HELB variants is measured on ssDNA-RPA tethers in channel 5. (D) and (E) Loop extrusion velocity and loop size for 10 nM HELB and 10 nM ΔHSM on human RPA-ssDNA; and of 10 nM HELB on yeast RPA-ssDNA. (F) Distribution of translocation processivity for 10 nM HELB, ΔOB, ΔHSM and ΔOBΔHSM on human RPA-ssDNA; and for 10 nM HELB on yeast RPA-ssDNA. For the interval scatter plots the center bar represents the mean of the data and the error bars represent the SD. Sample size is indicated below the data.

**Figure S5:**
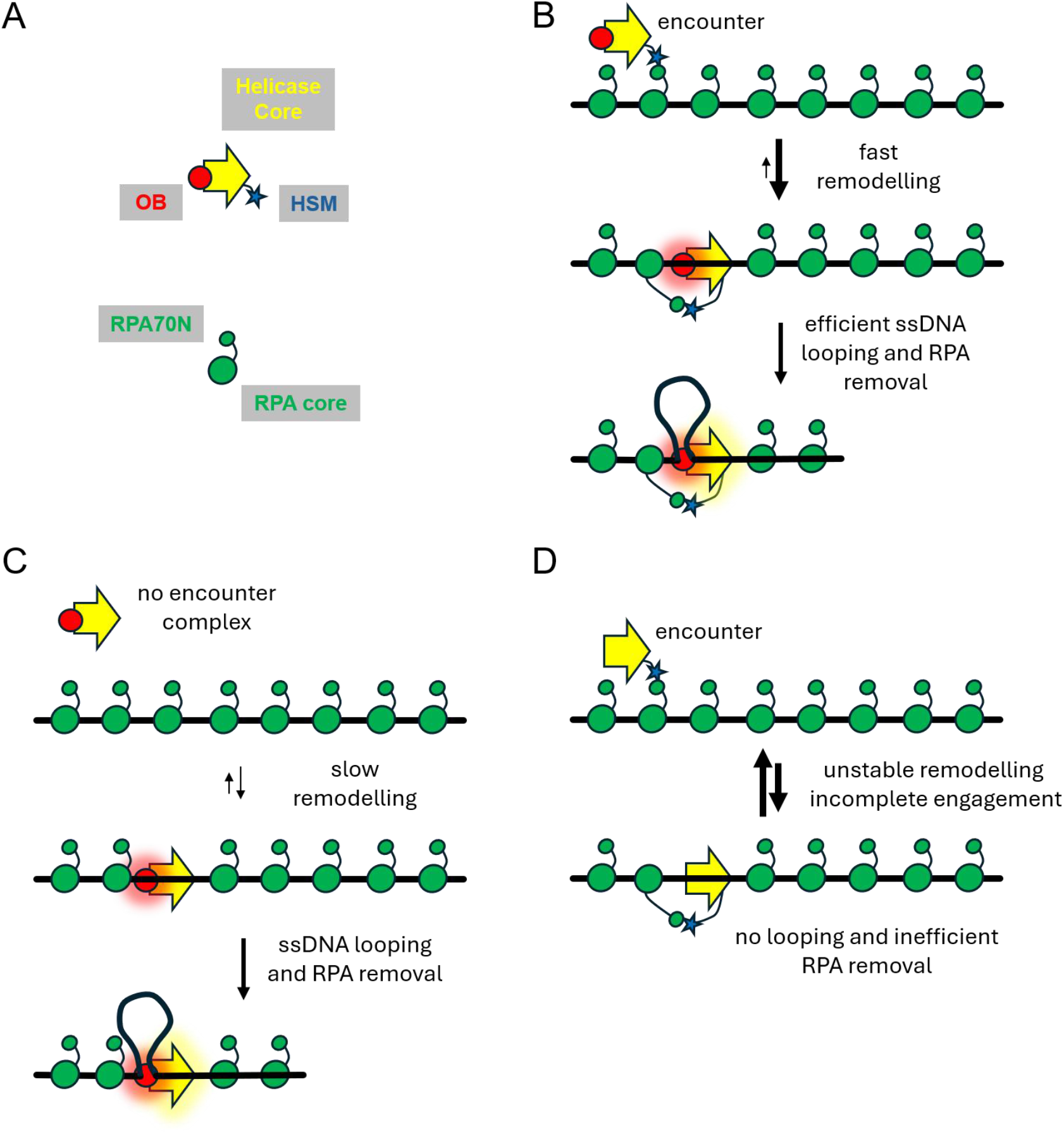
**A working model for the effects of OB and HSM domain deletion on HELB recruitment and activation at RPA-ssDNA filaments**. (A) Colour coding for domain organisation in cartoon models. (B) Schematic showing a working model for wild type HELB-dependent RPA removal. See also main text and main paper Figure 6 for further detail. (C) The ΔHSM variant is unable to form the initial encounter complex with the exterior of the RPA filament. Therefore, engagement of the core motor and OB anchor domains is slow and inefficient, limiting the overall process, but subsequent translocation, ssDNA looping and RPA removal (when it occurs) is essentially normal. (D) The ΔOB variant can form the initial encounter complex with the exterior of the RPA filament, but remodelling to engage the core motor domains is unstable due to the lack of the OB anchor domain. Failure to form ssDNA loops is associated with very poor RPA removal.

## REFERENCES

1. Hazeslip, L., et al., Genome Maintenance by DNA Helicase B. Genes (Basel), 2020. 11(5).

2. Singleton, M.R., M.S. Dillingham, and D.B. Wigley, Structure and mechanism of helicases and nucleic acid translocases. Annu Rev Biochem, 2007. 76: p. 23–50.

3. Tkac, J., et al., HELB Is a Feedback Inhibitor of DNA End Resection. Mol Cell, 2016. 61(3): p. 405–418.

4. Liu, H., P. Yan, and E. Fanning, Human DNA helicase B functions in cellular homologous recombination and stimulates Rad51-mediated 5’-3’ heteroduplex extension in vitro. PLoS One, 2015. 10(1): p. e0116852.

5. Taneja, P., et al., A dominant-negative mutant of human DNA helicase B blocks the onset of chromosomal DNA replication. J Biol Chem, 2002. 277(43): p. 40853–61.

6. Guler, G.D., et al., Human DNA helicase B (HDHB) binds to replication protein A and facilitates cellular recovery from replication stress. J Biol Chem, 2012. 287(9): p. 6469–81.

7. Guler, G.D., Z. Rosenwaks, and J. Gerhardt, Human DNA Helicase B as a Candidate for Unwinding Secondary CGG Repeat Structures at the Fragile X Mental Retardation Gene. Front Mol Neurosci, 2018. 11: p. 138.

8. Day, F.R., et al., Large-scale genomic analyses link reproductive aging to hypothalamic signaling, breast cancer susceptibility and BRCA1-mediated DNA repair. Nat Genet, 2015. 47(11): p. 1294–1303.

9. Stankovic, S., et al., Genetic links between ovarian ageing, cancer risk and de novo mutation rates. Nature, 2024. 633(8030): p. 608-614.

10. Ward, L.D., et al., Rare coding variants in DNA damage repair genes associated with timing of natural menopause. HGG Adv, 2022. 3(2): p. 100079.

11. Pan, Y., et al., Genetic variants in HELB contribute to premature ovarian insufficiency and early age of natural menopause. JCI Insight, 2025. 10(21).

12. Winship, A.L., et al., The importance of DNA repair for maintaining oocyte quality in response to anti-cancer treatments, environmental toxins and maternal ageing. Hum Reprod Update, 2018. 24(2): p. 119–134.

13. Chen, R. and M.S. Wold, Replication protein A: single-stranded DNA’s first responder: dynamic DNA-interactions allow replication protein A to direct single-strand DNA intermediates into different pathways for synthesis or repair. Bioessays, 2014. 36(12): p. 1156–61.

14. Caldwell, C.C. and M. Spies, Dynamic elements of replication protein A at the crossroads of DNA replication, recombination, and repair. Crit Rev Biochem Mol Biol, 2020. 55(5): p. 482–507.

15. Awate, S. and R.M. Brosh, Jr., Interactive Roles of DNA Helicases and Translocases with the Single-Stranded DNA Binding Protein RPA in Nucleic Acid Metabolism. Int J Mol Sci, 2017. 18(6).

16. Fanning, E., V. Klimovich, and A.R. Nager, A dynamic model for replication protein A (RPA) function in DNA processing pathways. Nucleic Acids Res, 2006. 34(15): p. 4126–37.

17. Pokhrel, N., et al., Dynamics and selective remodeling of the DNA-binding domains of RPA. Nat Struct Mol Biol, 2019. 26(2): p. 129–136.

18. Yates, L.A., et al., A structural and dynamic model for the assembly of Replication Protein A on single-stranded DNA. Nat Commun, 2018. 9(1): p. 5447.

19. Ahmad, F., et al., Hydrogen-deuterium exchange reveals a dynamic DNA-binding map of replication protein A. Nucleic Acids Res, 2021. 49(3): p. 1455–1469.

20. Arcus, V., OB-fold domains: a snapshot of the evolution of sequence, structure and function. Curr Opin Struct Biol, 2002. 12(6): p. 794–801.

21. Hormeno, S., et al., Human HELB is a processive motor protein that catalyzes RPA clearance from single-stranded DNA. Proc Natl Acad Sci U S A, 2022. 119(15): p. e2112376119.

22. Gilhooly, N.S., E.J. Gwynn, and M.S. Dillingham, Superfamily 1 helicases. Front Biosci (Schol Ed), 2013. 5(1): p. 206–16.

23. Tada, S., et al., Molecular cloning of a cDNA encoding mouse DNA helicase B, which has homology to Escherichia coli RecD protein, and identification of a mutation in the DNA helicase B from tsFT848 temperature-sensitive DNA replication mutant cells. Nucleic Acids Res, 2001. 29(18): p. 3835–40.

24. Gerhardt, J., G.D. Guler, and E. Fanning, Human DNA helicase B interacts with the replication initiation protein Cdc45 and facilitates Cdc45 binding onto chromatin. Exp Cell Res, 2015. 334(2): p. 283–93.

25. Wu, Y., et al., Structural characterization of human RPA70N association with DNA damage response proteins. Elife, 2023. 12.

26. Osei, B., et al., Rare SNP in the HELB gene interferes with RPA interaction and cellular function of HELB. NAR Mol Med, 2025. 2(2): p. ugaf019.

27. Abramson, J., et al., Accurate structure prediction of biomolecular interactions with AlphaFold 3. Nature, 2024. 630(8016): p. 493-500.

28. Dillingham, M.S., Superfamily I helicases as modular components of DNA-processing machines. Biochem Soc Trans, 2011. 39(2): p. 413–23.

29. Garcia, P.L., et al., RPA alleviates the inhibitory effect of vinylphosphonate internucleotide linkages on DNA unwinding by BLM and WRN helicases. Nucleic Acids Res, 2004. 32(12): p. 3771–8.

30. Nimonkar, A.V., et al., BLM-DNA2-RPA-MRN and EXO1-BLM-RPA-MRN constitute two DNA end resection machineries for human DNA break repair. Genes Dev, 2011. 25(4): p. 350–62.

31. Qin, Z., et al., Human RPA activates BLM’s bidirectional DNA unwinding from a nick. Elife, 2020. 9.

32. Gupta, R., et al., FANCJ (BACH1) helicase forms DNA damage inducible foci with replication protein A and interacts physically and functionally with the single-stranded DNA-binding protein. Blood, 2007. 110(7): p. 2390–8.

33. Suhasini, A.N., et al., FANCJ helicase uniquely senses oxidative base damage in either strand of duplex DNA and is stimulated by replication protein A to unwind the damaged DNA substrate in a strand-specific manner. J Biol Chem, 2009. 284(27): p. 18458–70.

34. Daley, J.M., et al., Multifaceted role of the Topo IIIalpha-RMI1-RMI2 complex and DNA2 in the BLM-dependent pathway of DNA break end resection. Nucleic Acids Res, 2014. 42(17): p. 11083–91.

35. Xue, C., et al., Single-molecule visualization of human BLM helicase as it acts upon double-and single-stranded DNA substrates. Nucleic Acids Res, 2019. 47(21): p. 11225–11237.

36. Stanage, T.H., et al., RPA exhaustion activates SLFN11 to eliminate cells with heightened replication stress. Nat Cell Biol, 2026.

37. Toledo, L.I., et al., ATR prohibits replication catastrophe by preventing global exhaustion of RPA. Cell, 2013. 155(5): p. 1088–103.

38. Fairhead, M. and M. Howarth, Site-specific biotinylation of purified proteins using BirA. Methods Mol Biol, 2015. 1266: p. 171–84.

39. Schindelin, J., et al., Fiji: an open-source platform for biological-image analysis. Nat Methods, 2012. 9(7): p. 676–82.

40. Aicart-Ramos, C., et al., Long DNA constructs to study helicases and nucleic acid translocases using optical tweezers. Methods Enzymol, 2022. 673: p. 311–358.

41. Crooks, G.E., et al., WebLogo: a sequence logo generator. Genome Res, 2004. 14(6): p. 1188–90.

42. Suzek, B.E., et al., UniRef: comprehensive and non-redundant UniProt reference clusters. Bioinformatics, 2007. 23(10): p. 1282–8.

43. Papadopoulos, J.S. and R. Agarwala, COBALT: constraint-based alignment tool for multiple protein sequences. Bioinformatics, 2007. 23(9): p. 1073–9.

